# B1-type cyclins control microtubule organization during cell division in Arabidopsis

**DOI:** 10.1101/2021.06.29.450310

**Authors:** Mariana R. Motta, Xin’Ai Zhao, Martine Pastuglia, Katia Belcram, Farshad Roodbarkelari, Maki Komaki, Hirofumi Harashima, Shinichiro Komaki, Petra Bulankova, Maren Heese, Karel Riha, David Bouchez, Arp Schnittger

## Abstract

Flowering plants contain a large number of cyclin families, each containing multiple members, most of which have not been characterized to date. Here, we analyzed the role of the B1 subclass of mitotic cyclins in cell cycle control during Arabidopsis development. While we reveal *CYCB1;5* to be a pseudogene, the remaining four members were found to be expressed in dividing cells. Mutant analyses showed a complex pattern of overlapping, development-specific requirements of B1-type cyclins with CYCB1;2 playing a central role. The double mutant *cycb1;1 cycb1;2* is severely compromised in growth, yet viable beyond the seedling stage, hence representing a unique opportunity to study the function of B1-type cyclin activity at the organismic level. Immunolocalization of microtubules in *cycb1;1 cycb1;2* and treating mutants with the microtubule drug oryzalin revealed a key role of B1-type cyclins in orchestrating mitotic microtubule networks. Subsequently, we identified the GAMMA-TUBULIN COMPLEX PROTEIN 3-INTERACING PROTEIN 1 (GIP1/MOZART) as an *in vitro* substrate of B1-type cyclin complexes and further genetic analyses support an important role in the regulation of GIP1 by CYCB1s.

## Introduction

A highly elaborated control system guides cells through mitosis during which chromosomes are separated and distributed to the newly forming daughter cells. Cyclin-dependent kinase (CDK)-cyclin complexes stand in the center of this control system (Morgan, 1997; Lindqvist *et al*, 2009). In animals, Cdk1 together with B-type cyclins are an essential part of the so-called mitosis promoting factor (MPF) complex that phosphorylates a plethora of mitotic substrates including nuclear structure proteins, such as Lamin A and B, and chromosome segregation proteins, such as the spindle assembly factor TPX2 (Blethrow *et al*, 2008). MPF activity is kept low prior to mitotic entry by excluding Cyclin B1 (CycB1) from the nucleus (Toyoshima *et al*, 1998; Yang *et al*, 1998; Hagting *et al*, 1998). In addition, Cdk1-cyclin B complexes are inhibited by phosphorylation on two inhibitory residues, Thr14 and Tyr15 (or the homologous amino acids in the P-loop of the respective Cdk) by the action of Wee1 and/or Myt1 kinases (O’Farrell, 2001). After a threshold concentration of Cdk1-CycB1 is reached, CycB1 accumulates in the nucleus and the Cdk-CycB1 complex becomes activated by a group of dual-specificity Cdc25 phosphatases that remove the inhibitory phosphorylation from the P-loop of the kinase. Due to a negative feedback wiring with Wee1 and a positive feedback with Cdc25 (Tyson & Novak, 2001), Cdk1-cyclin activity levels rise rapidly and promote entry and progression through mitosis, including the separation of the duplicated centrosomes (spindle pole body in yeast) as a key step to generate a bipolar spindle (Lacey *et al*, 1999; Haase *et al*, 2001). Finally, to complete mitosis and promote cytokinesis, Cdk1-cyclin B levels have to drop. This is accomplished by the degradation of cyclin B mediated by the Anaphase Promoting Complex/Cyclosome (APC/C) (Nakayama & Nakayama, 2005).

While a wealth of information about the execution of mitosis exists in animals and yeast, information is still scarce in plants. Notably, flowering plants appear to regulate mitosis differently from yeast and animals. First of all, flowering plants do not contain centrosomes and it is still not fully understood how the mitotic spindle is organized, although many microtubule-regulating components are conserved (Yamada & Goshima, 2017). Next, Arabidopsis WEE1 kinase was shown to prevent premature cell differentiation in S phase after DNA damage rather than functioning in mitotic control (De Schutter *et al*, 2007; Cools *et al*, 2011). Moreover, Arabidopsis does not contain a functional Cdc25 homolog, thus one of the most central control loops of the animal cell cycle is absent at least in this plant (Dissmeyer *et al*, 2010, 2009).

Another difference in mitotic regulation between plants and animals appears at the level of cyclins. In animals, D-type cyclins control entry into S-phase (G1 cyclins), while cyclin A controls S phase as well as early mitotic events, and B-type cyclins control mitosis (Furuno *et al*, 1999; Riabowol *et al*, 1989). In contrast, functional studies and expression analyses have revealed that members of all three cyclin classes, i.e., cyclin A, B, and D, are involved in the control of mitosis in plants (Schnittger *et al*, 2002; Boudolf *et al*, 2009; Menges *et al*, 2005; Dewitte *et al*, 2007; Vanneste *et al*, 2011). While there are only a few members in each cyclin family in metazoans, plant cyclin families are large, which makes functional studies challenging. For instance, as opposed to three B-type cyclins in mammals (CycB1, CycB2 and CycB3) and two in Drosophila (CycB1 and CycB3), there are eleven predicted B-type cyclins divided into three subgroups (B1, B2 and B3) in Arabidopsis that are all equally distant from animal B-type cyclins, i.e., Arabidopsis B1-type cyclins are closer related to B2 and B3 from Arabidopsis than to any B-type cyclin from animals (Vandepoele *et al*, 2002; Wang *et al*, 2004; Doerner *et al*, 1996). This classification is currently only based on sequence similarities and for B-type cyclins, as for most other cyclins in plants, the biological role is far from being understood.

Here, we present a functional analysis of the largest class of B-type cyclins in Arabidopsis, i.e., the five-member-containing B1 group. We reveal a central role for CYCB1;2 that is backed up by one or more of the other B1-type cyclins in a tissue-dependent manner. Unlike CycB1 mutants in mouse (Brandeis *et al*, 1998), Arabidopsis *cycb1;1 cycb1;2* double mutants are viable, presenting a unique opportunity to study cyclin B function at an organismic level. This allowed us to reveal the organization of mitotic microtubules as the main function of B1-type cyclins in Arabidopsis, a finding supported by *in vitro* kinase assays that indicated that GIP1/MOZART, a key factor of microtubule organization, is a substrate of CDK-CYCB1 complexes.

## Results

### CYCB1;1, CYCB1;2 and CYCB1;3 are redundantly required for endosperm proliferation

To start the characterization of B1-type cyclins, we first determined their expression pattern. To this end, we used previously generated promoter reporter lines comprising *GFP* fused to the N-terminal part of the respective cyclin (*CYCB1;1* to *CYCB1;4*), including the destruction box (Weimer *et al*, 2016). For *CYCB1;5*, since different annotations exist for this gene, we generated three different reporter constructs, each reaching to the different predicted transcriptional start sites. One upstream of the first ATG, one upstream of the second ATG and the third one including the first and second upstream regions. However, in none of these *CYCB1;5* reporter lines a signal could be detected. Therefore, we next analyzed the expression of *CYCB1;5* by qRT-PCR. Sequencing of the amplified products showed the existence of many different *CYCB1;5* cDNAs that exhibited exon skipping, intron retention and use of internal polyadenylation sites (Fig EV1), consistent with the lack of reliable transcriptional support for *CYCB1;5* in public depositories (The Arabidopsis Information Resource, TAIR). Taken together with data from previous studies (Bulankova *et al*, 2013) and the fact that many *Arabidopsis* accessions have accumulated several point mutations and even deletions in *CYCB1;5* (The Arabidopsis Information Resource, TAIR), we concluded that *CYCB1;5* is a pseudogene. In the following study, we therefore concentrated on the analysis of *CYCB1;1* through *CYCB1;4*.

The expression of the four B1-type cyclins has been previously described in a patchy pattern in regions with high cell proliferation activity, such as in the roots (Weimer *et al*, 2016). Hence, these cyclins seem to be true mitotically expressed genes. Furthermore, previous genome-wide expression studies have detected that the transcripts of all three B1-type cyclins *CYCB1;1*, *CYCB1;2* and *CYCB1;3* are overrepresented in the developing endosperm (Day *et al*, 2008). In agreement, we found that the promoter reporter constructs for *CYCB1;1*, *CYCB1;2* and *CYCB1;3* but not *CYCB1;4* were expressed during endosperm development (Fig EV2).

To assess the individual biological role of B1-type cyclins, we then analyzed previously isolated null mutants for all four B1-type cyclins (Weimer *et al*, 2016). However, none of the single mutants showed an obvious deviation from the wildtype under normal growth conditions, as for instance seen in growth and seed viability (Fig 1 and 2) in comparison to the wildtype. This finding was consistent with former observations (Weimer *et al*, 2016). Since *CYCB1;1*, *CYCB1;2* and *CYCB1;3* reporters have similar enrichment in the proliferating endosperm (Fig EV2), and *CYCB1;1*, *CYCB1;2, CYCB1;3* and *CYCB1;4* have a similar expression pattern in the roots, we reasoned that the B1-type cyclins might control mitotic divisions redundantly. Therefore, we also generated and analyzed all six possible double mutant combinations. The growth of the *cycb1;1 cycb1;2* double mutant was severely reduced (Fig 1B-D; for detailed characterization see below), while the size and morphology of the other double mutants were at a first look indistinguishable from the wildtype.

**Figure 1.**
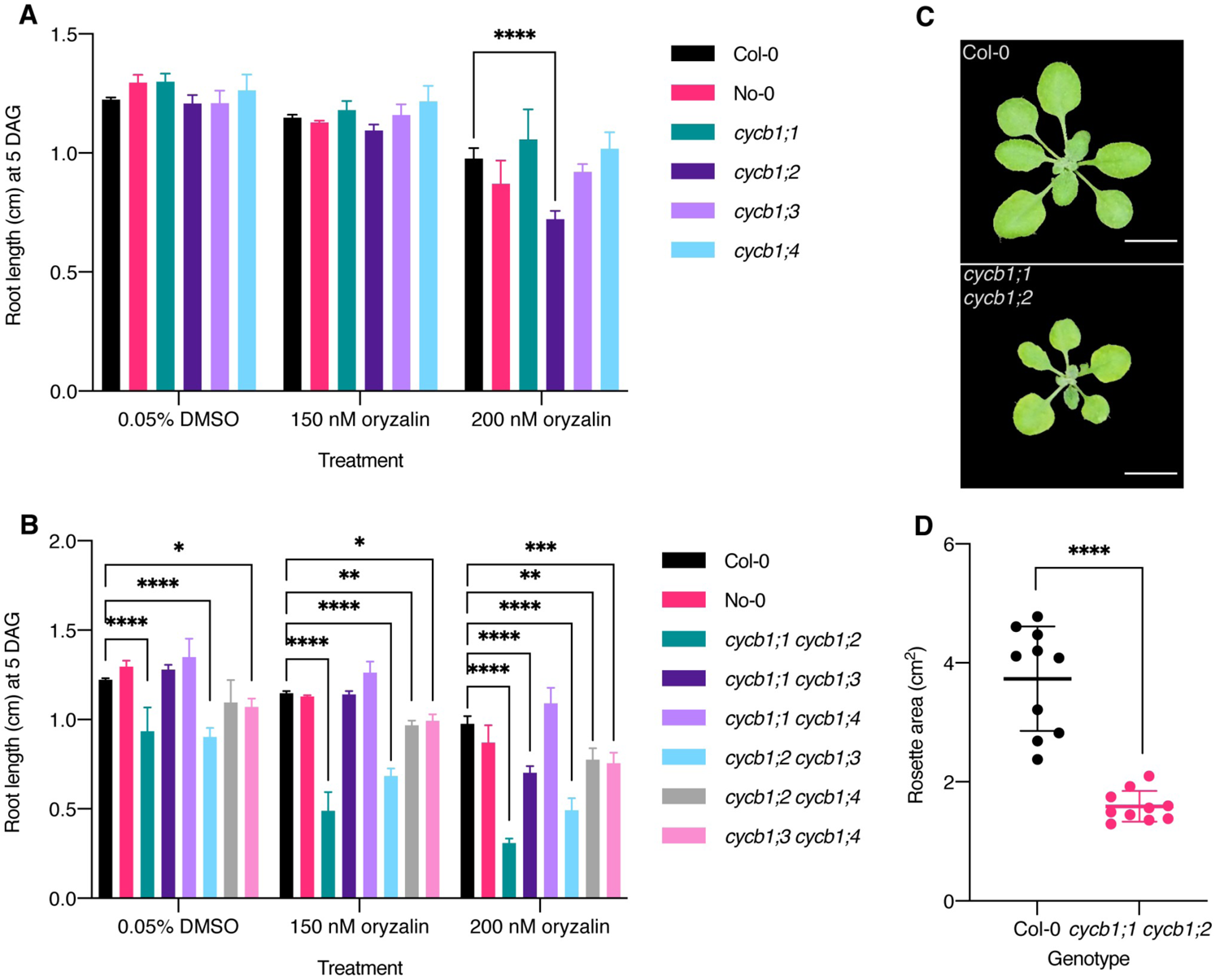
*cycb1;1 cyb1;2* mutants show decreased root growth and shoot development, which is further emphasized by a MT-destabilizing drug. A, B. Quantification of oryzalin root growth assays in single (A) and double (B) mutants. DAG = days after germination. Graphs show mean ± SD of three biological replicates with at least 10 plants per genotype per replicate. Asterisks indicate a significant difference in root length in a two-way ANOVA followed by Tukey’s multiple comparisons test. C. Rosette pictures of 20-day-old Col-0 and *cycb1;1 cycb1;2*. Scale bar: 1 cm. D. Quantification of the rosette area using total leaf surface in Col-0 and *cycb1;1 cycb1;2*. Graph represents the single rosette area values and the horizontal lines indicates the mean value ± SD, *n* = 10 plants per genotype. Asterisks indicate a significant difference in rosette area using an unpaired *t*-test, *t* = 7.421, df = 18.

**Figure 2.**
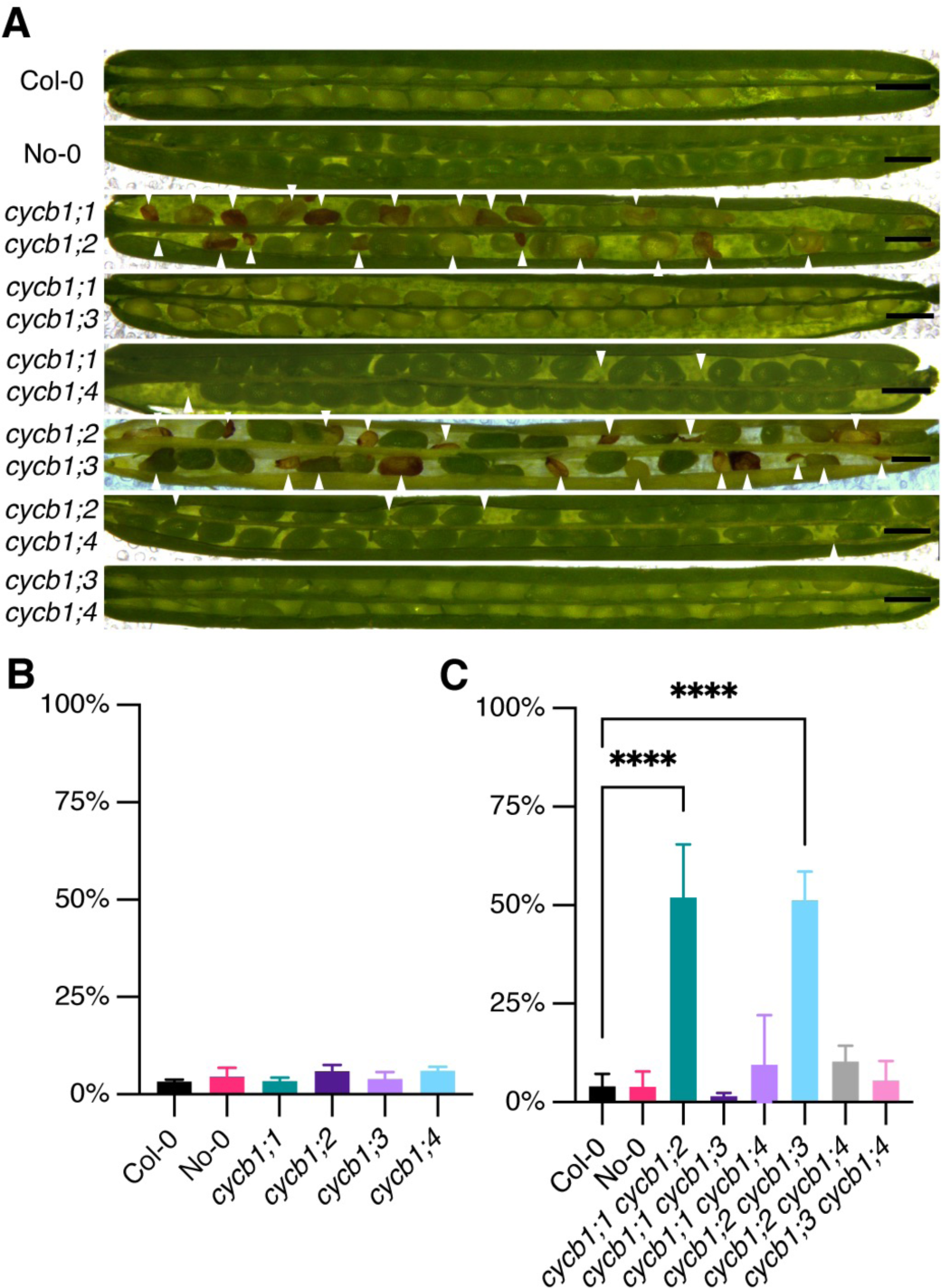
Seed abortion in *cycb1* mutants. A. Silique pictures of *cycb1* double mutant combinations. White arrowheads indicate aborted ovules and seeds. Scale bars: 500 μm. B, C. Quantification of aborted seeds in single (B) and double (C) mutants. Graphs represents the average seed abortion rate per plant ± SD of three biological replicates, *n* = 550-1029 seeds analyzed per genotype. Asterisks indicate significant differences in seed abortion rate in an ordinary one-way ANOVA test, followed by a Dunnett’s multiple comparisons test.

An analysis of the siliques in the double mutants revealed that *cycb1;1 cycb1;3*, *cycb1;1 cycb1;4*, and *cycb1;3 cycb1;4* did not have a reduced seed set. In contrast, *cycb1;1 cycb1;2* and *cycb1;2 cycb1;3* had on average approximately half of the seeds aborted (Fig 2A and 2C; 52.0% ± 13.5%, *n* = 3 biological replicates, 550 seeds, for *cycb1;1 cycb1;2* and 51.3% ± 7.2%, *n* = 3 biological replicates, 769 seeds, for *cycb1;2 cycb1;3* versus 4.0% ± 3.2%, *n* = 3 biological replicates, 821 seeds, for wildtype, Col-0; *P* < 0.0001 for both comparisons). The appearance of the aborted seeds varied in size and color (Fig 2A). Some seeds lacked the typical green color of a maturating embryo and looked transparent while others appeared brown and shriveled; in some cases, unfertilized or aborted ovules were visible.

To investigate the cause of this seed abortion, we collected, fixed and cleared seeds 3 days after pollination (DAP) (Fig 3). Since endosperm nuclei exhibit a strong autofluorescence, we were able to assess seed development quantitatively by using confocal laser scanning microscopy. In wildtype *Arabidopsis* seeds, a fertilized central cell will undergo seven to eight cycles of free nuclear divisions leading to an estimated total of more than 200 nuclei in the endosperm of three day old seeds (Boisnard-Lorig *et al*, 2001). In our analysis, the general morphology of the seeds at the single mutant level seemed unchanged in comparison to the wildtype (Fig 3A). However, counting the number of endosperm nuclei in these seeds 3 DAP displayed a strong reduction in endosperm divisions in the *cycb1;2* mutant (Fig 3B; 94.1 ± 55.4 endosperm nuclei per seed, *n* = 30) in comparison to the wildtype (Col-0, 175.2 ± 43.6, *n* = 30; *P* < 0.0001).

**Figure 3.**
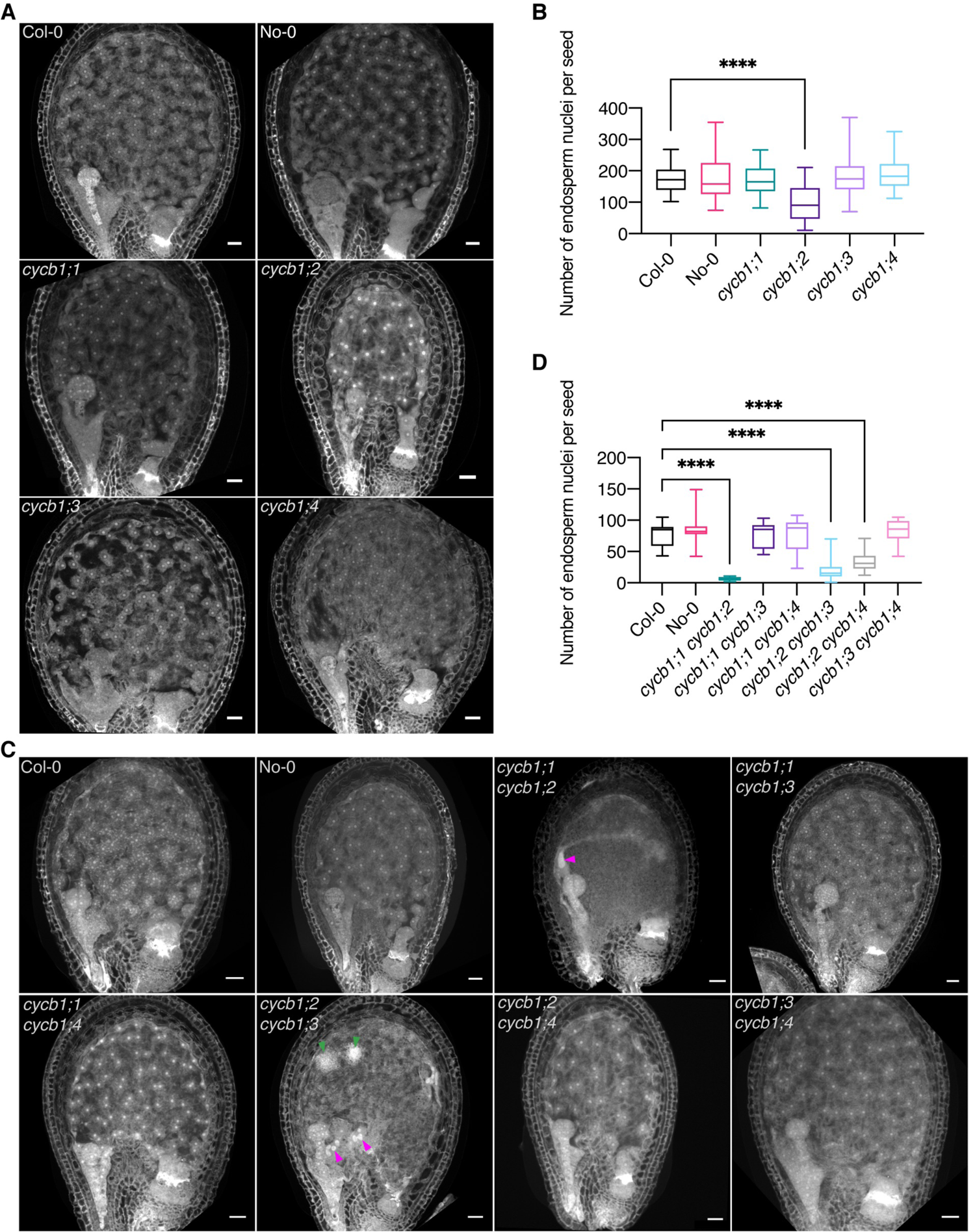
CYCB1 mutations delay endosperm proliferation. A, C. Confocal microscopy images of seeds 3 DAP. Endosperm and embryo morphology in *cycb1* single (A) and double (C) mutants. Magenta arrowheads indicate enlarged endosperm nuclei, while green arrowheads indicate atypical agglomerates of endosperm nuclei. Scale bars: 25 μm. B, D. Quantification of endosperm nuclei in *cycb1* single (B) and double (D) mutants. Boxes and whiskers represent min to max values with the median indicated as a central horizontal line, *n* = 26-30 seeds per genotype. Asterisks show significant differences in the number of endosperm nuclei per seed in a Kruskal-Wallis test followed by Dunn’s multiple comparisons test.

At the double mutant level, the general morphology of *cycb1;1 cycb1;2* and *cycb1;2 cycb1;3* seeds 3 DAP appeared abnormal (Fig 3C). The endosperm nuclei number at this time point was dramatically reduced with only 6 ± 3 in *cycb1;1 cycb1;2* (*n* = 26) and 28 ± 9 in *cycb1;2 cycb1;3* (*n* = 30) in relation to 77 ± 19.3 in the wildtype (Fig 3D; Col-0, *n* = 30; *P* < 0.0001 for both comparisons). Moreover, the nuclei appeared to be extremely enlarged in *cycb1;1 cycb1;2* and *cycb1;2 cycb1;3* (Fig 3C; magenta arrowheads), and in *cycb1;2 cycb1;3* atypical agglomerates of micro-sized nuclei were seen (Fig 3C; green arrowheads). The strong accumulation of a reporter gene when expressed from the *CYCB1;1*, *CYCB1;2* and *CYCB1;3* promoters in the seeds of these two double mutants consistently revealed enlarged nuclei which appeared, on the basis of reporter activity, to be halted at G2/M (Fig EV2B and EV2C). The double mutant *cycb1;2 cycb1;4* also showed a decrease in endosperm nuclei number (33.4 ± 12.87, *n* = 30) in relation to the wildtype (*P* < 0.0001), yet not as extensive as in the *cycb1;1 cycb1;2* and *cycb1;2 cycb1;3* double mutants and no major morphological abnormalities were identified, which could be explained by our observation that *CYCB1;4* was never expressed in the developing endosperm, consistent with the non-enrichment of the transcript in this tissue as shown in Day *et al*, 2008, and therefore CYCB1;4 might not play a major role in endosperm divisions.

Taken together, we conclude that CYCB1;2 is of major importance for the free nuclear divisions during endosperm development and acts redundantly with CYCB1;1 and CYCB1;3.

### CYCB1;1, CYCB1;2 and CYCB1;4 together control female gametophyte development

Up to this point, we did not find a clear role for CYCB1;4, suggesting even higher levels of redundancy among the B1-type cyclins or, alternatively, an overlapping function with other B-type cyclins in *Arabidopsis*. To clarify the relative contribution of the four CYCB1 genes to development, we decided to investigate the role of the CYCB1 group in detail by first constructing the triple *cycb1;1*^-/-^ *cycb1;3*^-/-^ *cycb1;4*^-/-^ mutant. Notably, this triple mutant was not different from the wildtype as for instance judged by overall growth, seed viability (Fig 4D), pollen development and pollen viability (Fig 5C and 5E). This finding further underlined the paramount role of CYCB1;2 among the B1-type cyclins. This result also indicated that CYCB1;4, if functionally relevant, may have a redundant role with either one or both of the two pairs CYCB1;1 CYCB1;2 and CYCB1;2 CYCB1;3. To test this, we generated the triple and quadruple mutant combinations *cycb1;1*^-/-^ *cycb1;2*^+/-^ *cycb1;4*^-/-^ and *cycb1;1^-/-^ cycb1;2^+/-^ cycb1;3*^+/-^ *cyb1;4*^-/-^. While overall growth of the triple and quadruple mutant combinations (note that at least one B1 gene is not homozygous mutant in these combinations) was similar to the wildtype, we found a strong reduction in fertility as siliques contained approximately 43% (± 0.4%, *n* = 3 biological replicates, 500 seeds; *P* < 0.0001) and 48% (± 2.1%, *n* = 3 biological replicates, 579 seeds; *P* < 0.0001) of aborting or unfertilized ovules and/or aborting seeds for *cycb1;1*^-/-^ *cycb1;2*^+/-^ *cycb1;4*^-/-^ and *cycb1;1^-/-^ cycb1;2^+/-^ cycb1;3*^+/-^ *cyb1;4*^-/-^ respectively in comparison to the wildtype (Fig 4D and 4E; 0.8% ± 0.9%, *n* = 3 biological replicates, 487 seeds). This abortion rate suggested a female gametophytic defect and we therefore analyzed embryo sac development in the mutants.

**Figure 4.**
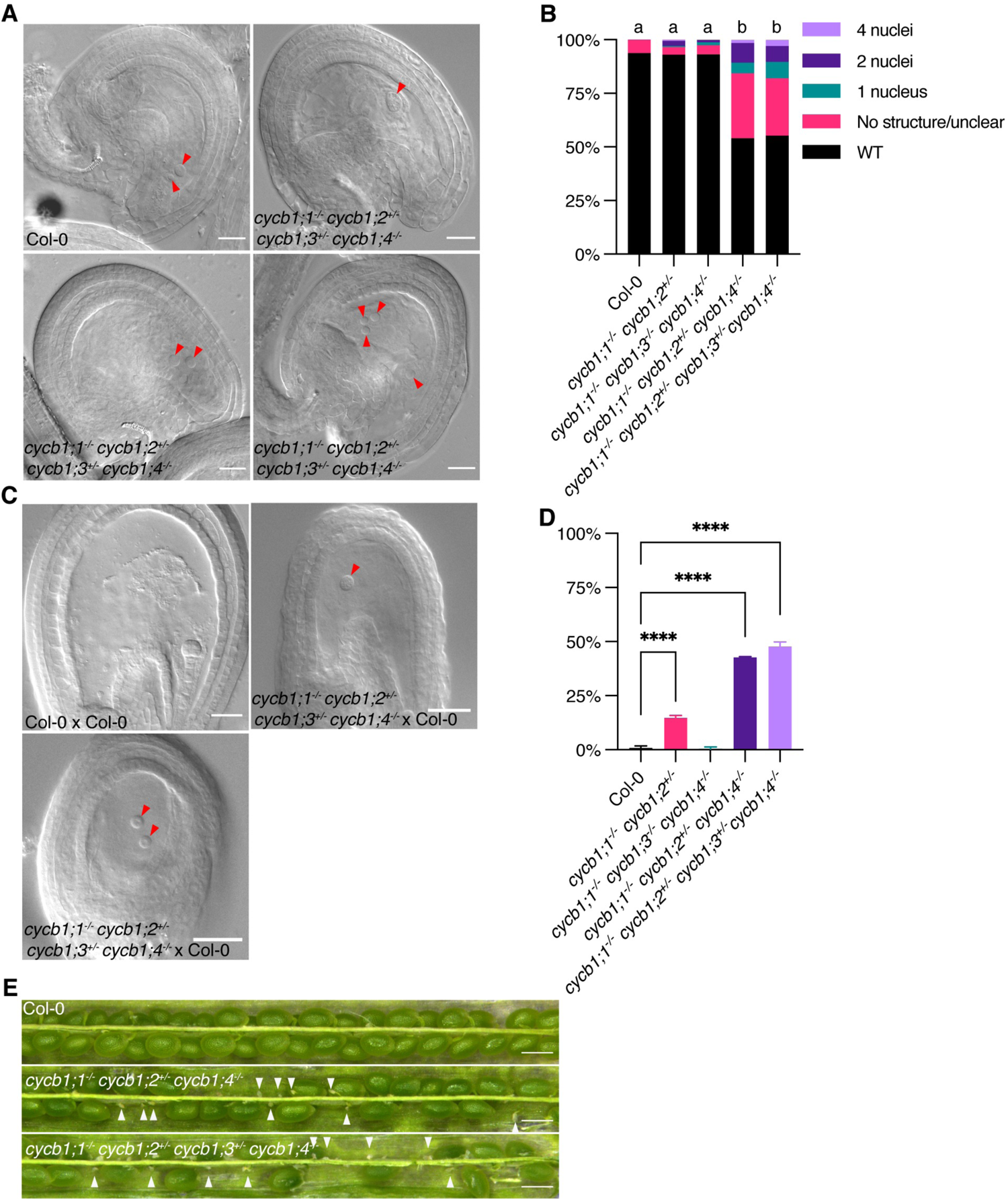
Embryo sac development is controlled by CYCB1 members. A. DIC images of abnormal embryo sacs in *cycb1* mutant combinations. Red arrowheads indicate the visible nuclei in Col-0 embryo sacs (central and egg cells) and the corresponding structures in the quadruple *cycb1;1^-/-^ cycb1;2^+/-^ cycb1;3*^+/-^ *cyb1;4*^-^ mutant. Scale bars: 20 μm. B. Quantification of the different abnormal embryo sac structures in *cycb1* mutant combinations (*n* = 202-459 embryo sacs per genotype). Different letters indicate significant differences in the proportion of abnormal embryo sacs in a Chi-squared test followed by the Marascuilo procedure to identify significant pairwise comparisons. WT = wildtype. C. DIC images of embryo sacs 3 DAP with wildtype pollen (female x male). Red arrowheads indicate the visible embryo sac nuclei in the crosses with the quadruple *cycb1;1^-/-^ cycb1;2^+/-^ cycb1;3*^+/-^ *cyb1;4*^-^ mutant as a female donor, while the control Col-0 x Col-0 cross exhibits a developing embryo. Scale bars: 20 μm. D. Quantification of seed abortion in different *cycb1* mutant combinations. Graph represents the average seed abortion rate per plant ± SD of three biological replicates, *n* = 463-579 seeds analyzed per genotype. Asterisks indicate significant differences in seed abortion rate in an ordinary one-way ANOVA test, followed by a Dunnett’s multiple comparisons test. E. Silique pictures of *cycb1* triple and quadruple mutants. White arrowheads indicate early aborted ovules. Scale bars: 500 μm.

**Figure 5.**
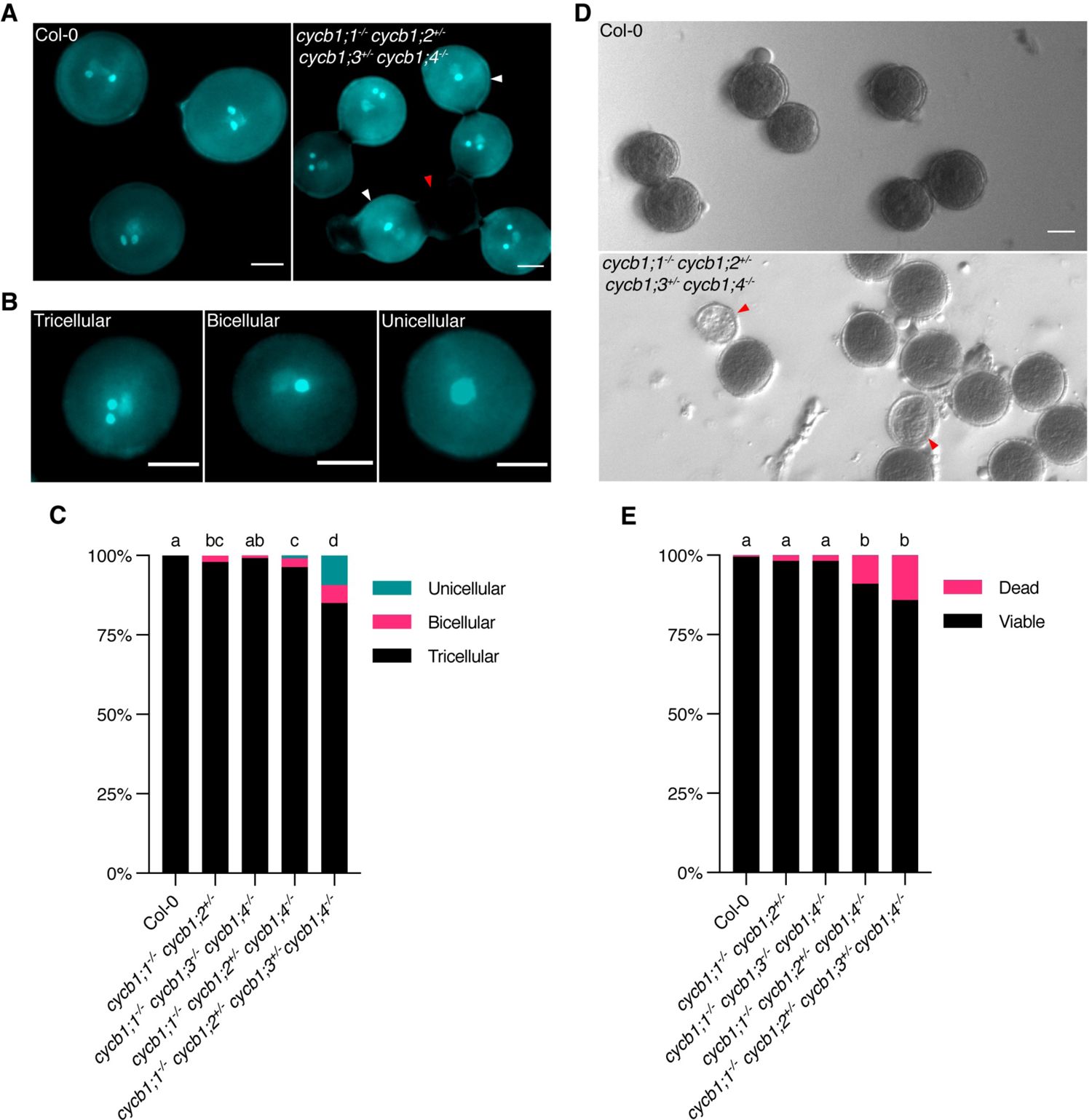
Pollen development is affected by mutations in the CYCB1 group. A, B. DAPI staining of pollen in *cycb1* mutants, including pollen configurations found in *cycb1;1*^-/-^ *cycb1;2*^+/-^ *cycb1;3*^+/-^ *cycb1;4*^-/-^ mutants (B). Scale bars: 5 μm. C. Quantification of DAPI-stained pollen configurations in different *cycb1* mutant combinations, *n* = 420-616 pollen grains per genotype. Different letters indicate significant differences in the proportion of abnormal pollen (uni- and bicellular) in a Chi-squared test followed by the Marascuilo procedure to identify significant pairwise comparisons. D. Alexander staining of mature pollen indicating pollen viability. Scale bars: 5 μm. E. Quantification of Alexander-stained pollen viability, *n* = 403-498 pollen grains per genotype. Different letters indicate significant differences in the proportion of dead pollen in a Chi-squared test followed by the Marascuilo procedure to identify significant pairwise comparisons. Red arrowheads indicate dead pollen, while white arrowheads indicate bicellular pollen.

In wild-type *Arabidopsis* plants, an embryo sac develops from a megaspore that is released after meiosis (Drews & Yadegari, 2002). Every megaspore undergoes three rounds of nuclear divisions resulting in an eight-celled embryo sac that subsequently cellularizes. The two centrally located polar nuclei fuse then to generate the central cell nucleus while the three antipodal cells that lay at the opposite side of the egg cell will undergo programmed cell death, resulting in a four-celled mature embryo sac that consists of a large, homodiploid central cell and an egg cell (red arrowheads; Fig 4A, Col-0) and two synergids that flank the egg cell (not shown). While this stereotypic wildtype developmental pattern was not significantly altered in *cycb1;1*^-/-^ *cycb1;2*^+/-^ double mutant combinations, consistent with the full transmission of the mutant allele through the female gametophyte (Table 1), we found embryo sacs from *cycb1;1*^-/-^ *cycb1;2*^+/-^ *cycb1;4*^-/-^ and *cycb1;1^-/-^ cycb1;2^+/-^ cycb1;3*^+/-^ *cyb1;4*^-/-^ mutant combinations with only one, two or four nuclei that did not show any sign of cellularization (Fig 4A); in addition, fuzzy embryo sacs were present in 30% and 27% of the cases respectively, likely indicating degenerating tissues, which is consistent with an early arrest of gametophytic development. In total, 46% (*n* = 459 embryo sacs analyzed) and 44.7% (*n* = 445) of embryo sacs from plants of the *cycb1;1*^-/-^ *cycb1;2*^+/-^ *cycb1;4*^-/-^ and *cycb1;1^-/-^ cycb1;2^+/-^ cycb1;3*^+/-^ *cyb1;4*^-/-^ combinations respectively were abnormal, in comparison to 6.2% (*n* = 210) in the wildtype (*P* < 0.0001) and 6.8% in the *cycb1;1*^-/-^ *cycb1;3*^-/-^ *cycb1;4*^-/-^ triple mutant (Fig 4B; P < 0.0001). The observation that the triple *cycb1;1*^-/-^ *cycb1;2*^+/-^ *cycb1;4*^-/-^ and quadruple *cycb1;1^-/-^ cycb1;2^+/-^ cycb1;3*^+/-^ *cyb1;4*^-/-^ mutants displayed a similar number of mutant embryo sacs (Fig 4B) suggests that CYCB1;3 is not required, at least at this triple mutant level, for the divisions of the female gametophyte.

**Table 1.** Transmission efficiency of the cycb1;1, cycb1;2 and cycb1;3 mutant alleles in reciprocal crosses of a triple mutant with wildtype. A transmission efficiency of 100% indicates full transmission of the mutant allele, i.e. 50% of the genotyped F1 seedlings are heterozygous. A z-test for one proportion was performed to test if the observed transmission frequencies differ from expected values and the significance level was corrected for multiple comparisons using Bonferroni. Asterisks indicate significant differences from expected values. Primers used for genotyping are listed in Table S1.

| Parental genotypes<br>(female x male) | Mutant allele |  |  | Number of<br>seeds (*n*) |
| --- | --- | --- | --- | --- |
|  | *cycb1;1* | *cycb1;2* | *cycb1;3* |  |
| *cycb1;1*<sup>+/-</sup> *cycb1;2*<sup>+/-</sup> *cycb1;3*<sup>+/-</sup> x Col-0 | 100% | 103% | 109.2% | 192 |
| Col-0 x *cycb1;1*<sup>+/-</sup> *cycb1;2*<sup>+/-</sup> *cycb1;3*<sup>+/-</sup> | 86%**** | 78.88%**** | 81.1%**** | 360 |

To assess the functionality of these embryo sacs, we pollinated the quadruple mutant combination with pollen from wildtype plants (Fig 4C). Supporting a female gametophytic defect, we observed a similar proportion of unfertilized and/or arrested embryo sacs in these crosses at 3 DAP (44.3%, *n* = 684 seeds) in comparison to control crosses in which wildtype plants were used as a female parent fertilized with wildtype pollen (0.7% arrested embryo sacs, *n* = 597). In reciprocal control crosses with pollen from mutant plants onto stigmas of wildtype plants, embryo and endosperm were formed in the developing seeds (seed abortion = 2.9%, *n* = 902 seeds). Thus, CYCB1;4, next to CYCB1;1 and CYCB1;2 appears to be required for embryo sac development. This was corroborated by analyzing the transmission of the *cycb1;2* and *cycb1;3* mutant alleles in reciprocal crosses of wildtype plants with the quadruple *cycb1;1*^-/-^ *cycb1;2^+/-^ cycb1;3^+/-^ cycb1;4^-/-^* mutant. As expected, transmission of *cycb1;2* was abolished through the female gametophyte (0%) and the efficiency in transmission was clearly reduced through the male gametophyte (70.8%) (Table 2), while the *cycb1;3* allele could be transmitted without an obvious reduction in efficiency through the females (92.7%).

**Table 2.** Transmission efficiency of the *cycb1;2* and *cycb1;3* mutant alleles in reciprocal crosses of a quadruple mutant with wildtype. A transmission efficiency of 100% indicates full transmission of the mutant allele, i.e. 50% of the genotyped F1 seedlings are heterozygous. A z-test for one proportion was performed to test if the observed transmission frequencies differ from expected values and the significance level was corrected for multiple comparisons using Bonferroni. Asterisks indicate significant differences from expected values. Primers used for genotyping are listed in Table S1.

| Parental genotypes<br>(female x male) | Mutant allele |  | Number of seeds<br>(*n*) |
| --- | --- | --- | --- |
|  | *cycb1;2* | *cycb1;3* |  |
| *cycb1;1<sup>-/-</sup>* *cycb1;2<sup>+/-</sup>* *cycb1;3<sup>+/-</sup>* *cycb1;4<sup>-/-</sup>*<br>x Col-0 | 0%**** | 92.7%**** | 192 |
| Col-0 x *cycb1;1<sup>-/-</sup>* *cycb1;2<sup>+/-</sup>* *cycb1;3<sup>+/-</sup>* *cycb1;4<sup>-/-</sup>* | 70.84%**** | 30.2%**** | 192 |

Interestingly, the requirement of the B1-type cyclins was different on the male side. Pollen develops from microspores through two consecutive divisions resulting in a tricellular grain that harbors two sperms next to one vegetative cell (McCormick, 2004) (Fig 5). We observed that both the *cycb1;2* as well as the *cycb1;3* mutant alleles were not fully transmitted through pollen in a cross of *cycb1;1*^-/-^ *cycb1;2^+/-^ cycb1;3^+/-^ cycb1;4^-/-^* with wildtype plants (Table 2; 70.8% and 30.2% transmission efficiency, respectively), indicating that all four B1-type cyclins contribute to the mitotic divisions of the developing pollen grain. Consistent with the reduced transmission, we also found pollen grains in mature anthers of *cycb1;1*^-/-^ *cycb1;2*^+/-^ *cycb1;4*^-/-^ and *cycb1;1^-/-^ cycb1;2^+/-^ cycb1;3*^+/-^ *cyb1;4*^-/-^ mutant combinations that comprised instead of three, only two or sometimes even one cell (Fig 5A-C). Accordingly, differential staining of aborted and non-aborted pollen showed an increased pollen abortion in the triple and quadruple mutants (Fig 5D and 5E) to 8.9% (*n* = 403 pollen grains analyzed) and 14.1% (*n* = 467), respectively, in relation to the wildtype (Col-0, 0.5%, *n* = 404; *P* < 0.0001).

Taken together, CYCB1;2 is also the most important B1 type cyclin during gametophyte development. CYCB1;3 appears to have only a minor role during female gametophyte development where instead CYCB1;4 acts together with CYCB1;1 and CYCB1;2. Remarkably, after fertilization the requirement changes, as presented above, and CYCB1;3 instead of CYCB1;4 is necessary for endosperm development.

### Root growth under microtubule-destabilizing conditions underlines the redundant role of CYCB1;1, CYCB1;2 and CYCB1;3 in regulating the cytoskeleton

Considering the severe reduction of growth in *cycb1;1 cycb1;2* mutants (Fig 1; see above) and to explore the role of CYCB1;2 and the other B1-type cyclins in controlling the microtubule cytoskeleton, we next analyzed the growth of B1-type cyclin mutants on medium containing the microtubule poison oryzalin. The rationale is that a minor defect in the mutants could become more prominent if the microtubule cytoskeleton is already slightly compromised. To that end, we compared the root growth of *cycb1* single and double mutants in ½ MS medium containing 150 nM or 200 nM oryzalin (Fig 1A and 1B respectively; Fig EV3 and EV4). As shown above, under control conditions (0.05% DMSO), the single *cycb1* mutants had similar root growth to the wildtype (Fig 1A). Once oryzalin was applied at 200 nM, *cycb1;2* grew significantly less (0.7 ± 0.03 cm, *n* = 3 biological replicates with at least 10 plants each) when compared to the wildtype (Col-0, 1.0 ± 0.04 cm, *n* = 3; *P* < 0.0001).

At the double mutant level, some combinations already showed a significantly shorter root even in control conditions (Fig 1B), such as *cycb1;1 cycb1;2* (0.9 ± 0.1 cm, *n* = 3; *P* < 0.0001) and *cycb1;2 cycb1;3* (0.9 ± 0.05 cm, *n* = 3; *P* < 0.0001) in comparison to the wildtype (Col-0, 1.2 ± 0.009 cm, *n* = 3). When 150 nM oryzalin was applied, the growth of Col-0 was reduced by approximately 6%, while *cycb1;1 cycb1;2* had a reduction of almost 50% in growth and *cycb1;2 cycb1;3* had a reduction of around 24%. As the concentration of oryzalin increased to 200 nM oryzalin, the difference in growth between Col-0 and other double mutants, such as *cycb1;2 cycb1;4* and *cycb1;3 cycb1;4*, became more pronounced. However, most strikingly, *cycb1;1 cycb1;3* mutants, which so far had shown no specific phenotype and no reduction in shoot or root growth, grew significantly shorter at 200 nM oryzalin (0.7 ± 0.04 cm, *n* = 3) in comparison to the wildtype (Col-0, 1.0 ± 0.04 cm, *n* = 3; *P* < 0.0001). Collectively, a major function of all four B1-type cyclins seems to be regulating the microtubule cytoskeleton.

### CYCB1;1 and CYCB1;2 control the organization of different microtubule arrays in the roots

Following our finding that *cycb1;1 cycb1;2* and *cycb1;2 cycb1;3* mutants have a severe reduction in root growth both in control and especially in microtubule-depolymerizing conditions, we performed whole mount immunolocalization studies against α-tubulin and KNOLLE, which is a G2/M and cell plate marker (Fig 6 and Fig 7; Table EV1). By analyzing mitotic divisions in the roots of these double mutants, a more detailed picture of cell division appeared.

**Figure 6.**
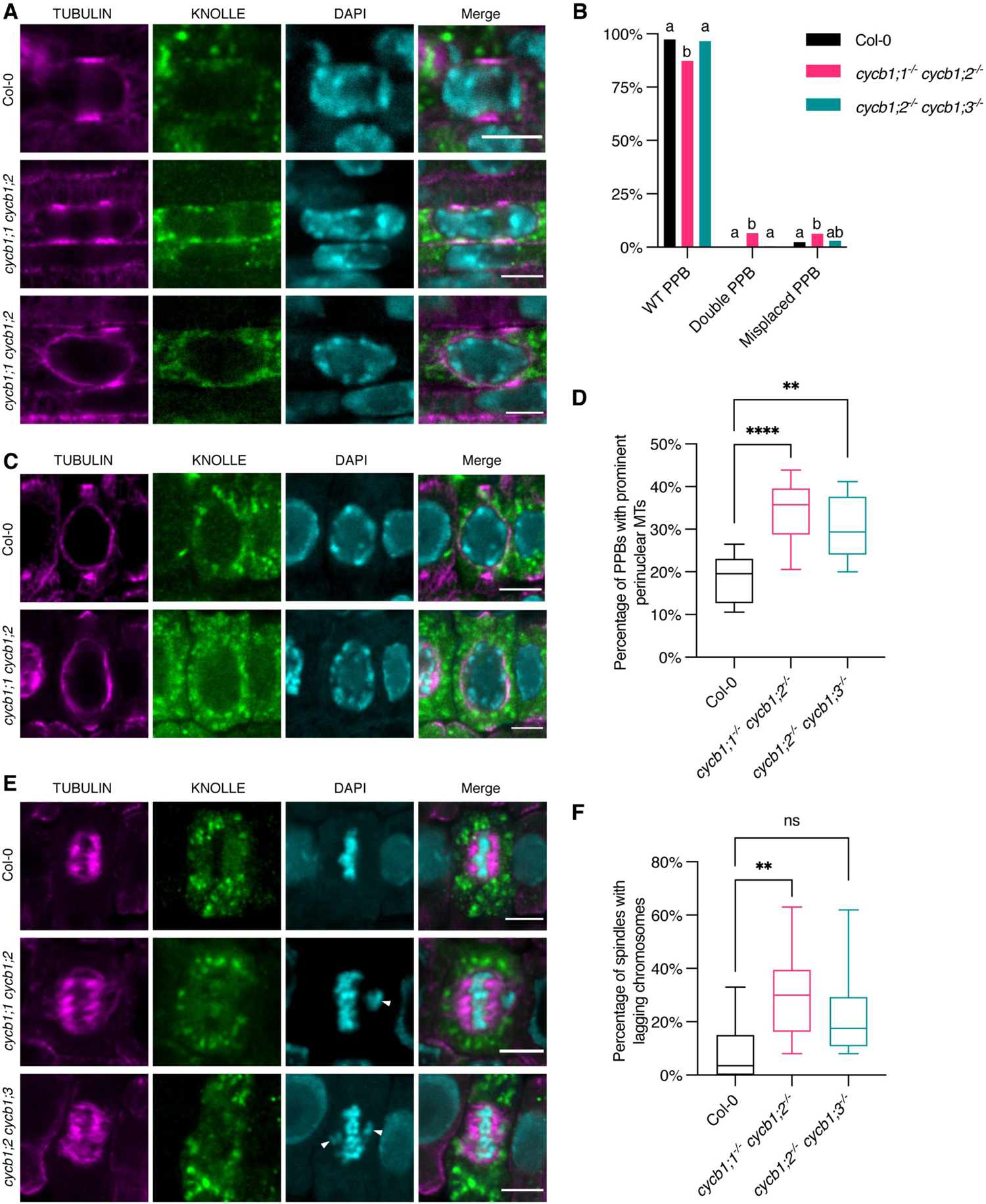
The double *cycb1;1 cycb1;2* mutant has abnormal microtubule arrays. A, C, E. Co-immunolocalization against tubulin (magenta) and KNOLLE (green) in root meristematic cells. Nuclei were counterstained with DAPI for the DNA (cyan). White arrowheads indicate laggards in the metaphase stage. Scale bars: 5 μm. B. Quantification of wildtype (WT), double and misplaced PPBs. Different letters indicate significant differences in the proportions of the different arrays per category in a Chi-squared test followed by the Marascuilo procedure to identify significant pairwise comparisons. Ten roots were analyzed per genotype. D. Quantification of PPBs with prominent perinuclear microtubules (MTs). Boxes and whiskers represent min to max values with the median indicated as a central horizontal line, *n* = 10 roots per genotype. Asterisks show significant differences in the percentage of PPBs with prominent perinuclear microtubules per root in an ANOVA test followed by Dunnett’s multiple comparisons test. F. Quantification of spindles with lagging chromosomes. Boxes and whiskers represent min to max values with the median indicated as a central horizontal line, *n* = 10 roots per genotype. Asterisks show significant differences in the percentage of spindles with lagging chromosomes per root in a Kruskal-Wallis test followed by Dunn’s multiple comparisons test.

**Figure 7.**
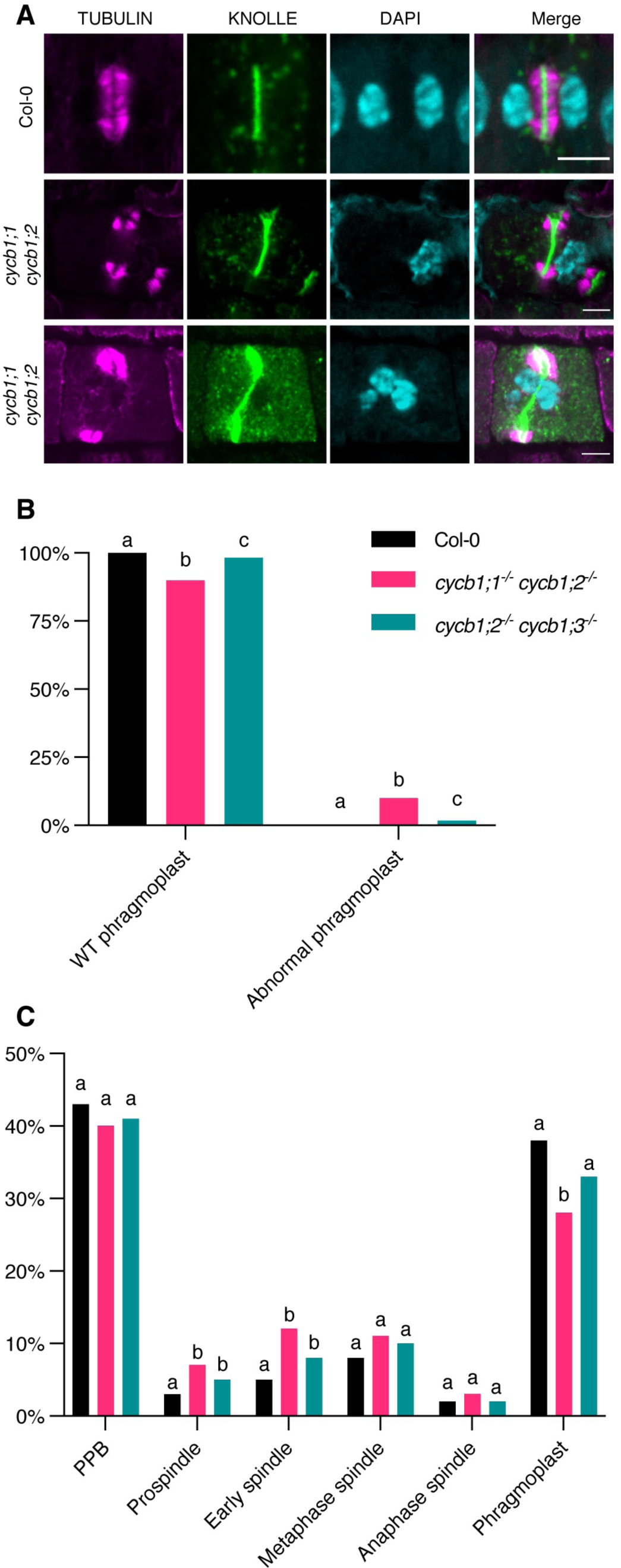
Both *cycb1;1 cycb1;2* and *cycb1;2 cycb1;3* have abnormal phragmoplasts and extended spindle stages. A. Co-immunolocalization against tubulin (magenta) and KNOLLE (green) in root meristematic cells. Nuclei were counterstained with DAPI for the DNA (cyan). Scale bars: 5 μm. B. Quantification of wildtype (WT) and abnormal phragmoplasts. Different letters indicate significant differences in the proportions of the different arrays per category in a Chi-squared test followed by the Marascuilo procedure to identify significant pairwise comparisons. Ten roots were analyzed per genotype. C. Quantification of the different mitotic stages in roots of the different genotypes. Different letters indicate significant differences in the proportions of the different arrays per category in a Chi-squared test followed by the Marascuilo procedure to identify significant pairwise comparisons. Ten roots were analyzed per genotype.

Preceding a mitotic division, a band of microtubules that encircles the nucleus at the equatorial plane, the so-called preprophase band (PPB), will form at the site of the future division plane. The PPB functions as a positional cue and anchoring site for proteins involved in the cell division site determination and by that contributes to the robustness of cell divisions (Schaefer *et al*, 2017). Following nuclear envelope breakdown, the barrel-shaped acentrosomal spindle that is responsible for separating the chromosomes forms. After proper bipolar kinetochore-microtubule attachments occur and enough tension is sensed, sister chromatids are pulled towards opposing poles. Next, the phragmoplast, which is a bipolar microtubule structure that expands in time towards the cell cortex, forms. The phragmoplast is a scaffold for cell wall formation on which vesicles are transported towards the microtubule-devoid midzone, where a growing cell plate is located.

As the cell progresses from G2 towards mitosis, the nuclear surface becomes a prominent site of microtubule nucleation. In *cycb1;1 cycb1;2* and *cycb1;2 cycb1;3* mutants, at the PPB stage, we observed an increase in perinuclear microtubules, with an average of 18.3% PPBs with prominent microtubules in Col-0 versus 34% and 30% in *cycb1;1 cycb1;2* and *cycb1;2 cycb1;3* respectively (Fig 6C and 6D). Either *cycb1* mutations induce an early accumulation of perinuclear microtubules or else the “mature” PPB stage, i.e. with perinuclear microtubules, lasts longer in the *cycb1* mutants, and cells have trouble progressing into mitosis. In the stele and pericycle cells of the *cycb1;1 cycb1;2* double mutant, we also observed cells harbouring double PPBs and cells with misplaced PPBs, i.e. PPBs that did not align properly at the equatorial plane of the nucleus. These double and misplaced PPBs were rarely seen in Col-0 wildtype plants; we found 6.50% double PPBs in *cycb1;1 cycb1;2* in comparison to 0.22% in wildtype (P < 0.0001) and 6.21% misplaced PPBs in comparison to 2.38% in wildtype (P = 0.009; Fig 6A and 6B).

At the spindle stage, irregular chromosome configurations were observed in *cycb1;1 cycb1;2*, with a significantly larger number of metaphase and anaphase spindles with chromosome laggards (Fig 6E and 6F). Although the number of spindles with lagging chromosomes in *cycb1;2 cycb1;3* was not significantly larger than in the wildtype, the impairment seen in those mutants in chromosome alignment was much more severe than that of the wildtype plants, i.e. chromosomes were seen far away from the metaphase plate. Finally, abnormal phragmoplasts were observed in the two double mutant combinations, including fragmented phragmoplasts, deformed phragmoplasts around abnormally shaped nuclei, and daughter cells with incompletely separated nuclei (*P* < 0.0001; Fig 7A and 7B). These abnormal phragmoplasts were likely a consequence of the irregular chromosome alignment and segregation seen in metaphase and anaphase. In short, all microtubule arrays were affected in the double mutants to a smaller or larger degree.

Next, we analyzed the proportion of cells at PPB, spindle and phragmoplast stages per root (Fig 7C). A significantly larger proportion of cells in both prospindle and early spindle stages were observed in *cycb1;1 cycb1;2* and *cycb1;2 cycb1;3* mutants (*P* < 0.0001), which indicates that these stages are delayed in those mutants. The phragmoplast stage is proportionally shorter in the *cycb1;1 cycb1;2* mutant, although this could be a direct consequence of the extended spindle stage since, if the proportions of some stages increase, the others decrease automatically. Accordingly, a flow cytometrical analysis revealed that *cycb1;1 cycb1;2* mutants have a higher proportion of 4C, 8C and 16C nuclei in comparison to the wildtype (Fig EV5), which is an indication that these mutants have longer G2 and/or M phases. An increase in polyploid cells could have two, not mutually excluding reasons. First, a failure to undergo cytokinesis. Second, a compromised division program leading to premature exit from proliferation and entry into differentiation accompanied by endoreplication. In addition, broader peaks were observed, suggesting the formation of aneuploidies as result of irregular mitotic divisions in this genotype.

In summary, CYCB1;1 and CYCB1;2 seem to be both redundantly required for robust root mitotic divisions under normal conditions, with CYCB1;3 playing a secondary role.

### The CYCB1 group forms active complexes mainly together with CDKB2;2 and can phosphorylate a MAP

Previous studies have shown that all B1-type cyclins can interact with all five major cell-cycle CDKs from *Arabidopsis*, i.e. CDKA;1, CDKB1;1, CDKB1;2, CDKB2;1, and CDKB2;2 (Van Leene *et al*, 2010). However, when we assessed the biochemical activity of all four B1-type cyclins with CDKA;1, CDKB1;1 and CDKB2;2, as representative members of the major cell-cycle CDKs, in comparative *in vitro* kinase assays against Histone H1, a more complex pattern appeared (Fig 8A). As a general principle, all four B1-type cyclins build the most active complexes with CDKB2;2, which is strictly expressed in mitosis. CYCB1;1 and CYCB1;4 showed overall the highest activity levels with CDKB2;2, followed by CYCB1;2 with CDKB2;2, while the CYCB1;3-CDKB2;2 pair was the least active among the CYCB1-CDKB2 complexes. Although much less than in complex with CDKB2;2, CYCB1;1, CYCB1;2 and CYCB1;4 could also phosphorylate Histone H1 together with CDKB1;1, but little to no activity was found in complexes with CDKA;1. In contrast, we could not detect any activity of CYCB1;3-CDKB1;1 complexes, while CYCB1;3 with CDKA;1 was almost as active as CYCB1;3-CDKB2;2 pairs.

**Figure 8.**
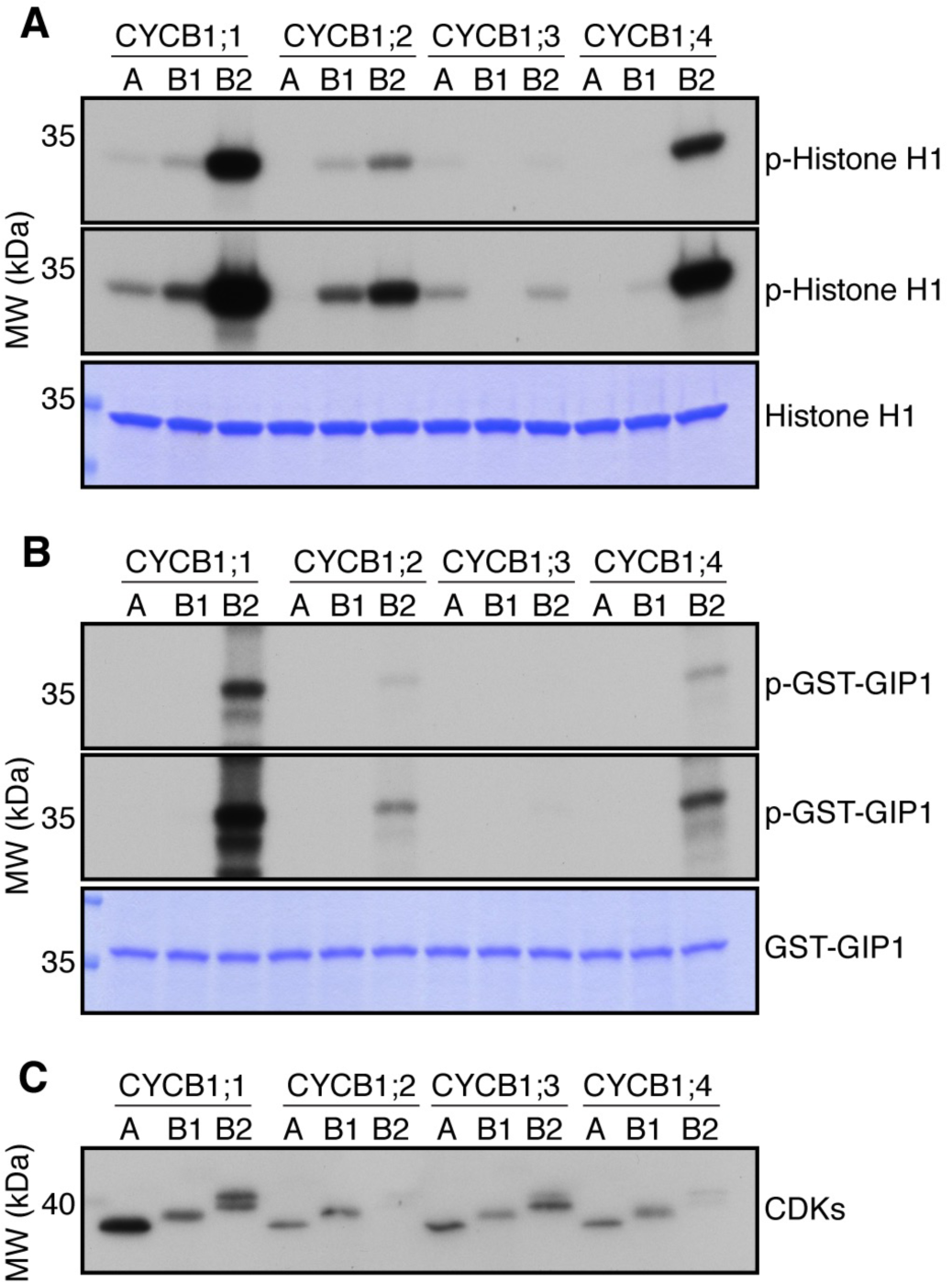
CYCB1-CDKB2;2 complexes are the most active and are able to phosphorylate a MT-nucleation factor. A. Kinase assays against Histone H1. Top and middle panels indicate shorter and longer exposures respectively of the same kinase assays. Bottom panel is a CBB staining of Histone H1 showing equal loading of the protein. A: CDKA;1, B1: CDKB1;1, B2: CDKB2;2. B. Kinase assays against GIP1. Top and and middle panels indicate shorter and longer exposures respectively of the same kinase assays. Bottom panel is a CBB staining of GIP1 showing equal loading of the protein. A: CDKA;1, B1: CDKB1;1, B2: CDKB2;2. C. Western blot against StrepIII-tagged proteins to show loaded amounts of the CDKs. A: CDKA;1, B1: CDKB1;1, B2: CDKB2;2.

The abnormal microtubule pattern observed in *cycb1;1 cycb1;2* mutants was reminiscent of the defects observed in microtubule binding and organizing protein mutants, such as in *gip1 gip2* double mutants (Janski *et al*, 2012; Nakamura *et al*, 2012), which are homologs of MOZART1 in animals. The *gip1 gip2* double knockdown mutant displays growth defects, sterility, defective microtubule arrays and spindles with irregular polarity, which is linked to chromosome laggards in metaphase and anaphase and aneuploidy (Janski *et al*, 2012). Additionally, the γ-tubulin *tubg1 tubg2* mutants display similar aberrant female and male gametophytes, with abnormal embryo sacs and reduced pollen nuclei number (Pastuglia *et al*, 2006). This suggested that CDK complexes containing B1-type cyclins might phosphorylate the GIPs and/or other microtubule organizing proteins. Indeed, GIP1 but not GIP2 contains a consensus CDK phosphorylation site at position T67. Therefore, we expressed GIP1 in bacteria and subjected it to *in vitro* kinase assays with all four CYCB1 members each paired with either CDKA;1, CDKB1;1 or CDKB2;2. High activity levels against GIP1 were found for CYCB1;1, CYCB1;2 and CYCB1;4 (Fig 8B). However, these B1-type cyclins phosphorylated GIP1 only in combination with CDKB2;2, highlighting the importance of both the cyclin and the CDK partner for substrate recognition in plants and further emphasizing B2-type CDKs as the most important partners of the cyclin B1 group.

Following the finding that GIP1 is phosphorylated by CYCB1-CDKB2;2 complexes, we decided to generate triple *gip1 cycb1;1 cycb1;2* and *gip2 cycb1;1 cycb1;2* mutants. However, we were never able to isolate *gip2 cycb1;1 cycb1;2* mutants (Table 3). To address whether the missing triple mutant was due to a gametophytic and/or embryonic defect, we performed reciprocal crosses with *gip1*^-/-^ *cycb1;1^-/-^ cycb1;2^+/-^* and *gip2*^-/-^ *cycb1;1^-/-^ cycb1;2^+/-^* as male and female donors with Col-0 (Table 4). With the exception of a reduced transmission efficiency of *cycb1;2* through the female gametophyte of approximately 65% in *gip2*^-/-^ *cycb1;1^-/-^ cycb1;2^+/-^* crosses with the wildtype, we observed that both *gip1 cycb1;1 cycb1;2* and *gip2 cycb1;1 cycb1;2* gametes were largely viable and transmitted both through the female and male sides. This indicated that the triple *gip2*^-/-^ *cycb1;1^-/-^ cycb1;2^-/-^* mutation is embryo lethal.

**Table 3.** Distortion of cycb1;2 segregation in a gip2^-/-^ cycb1;1^-/-^ cycb1;2^+/-^ background. The expected mendelian segregation reflects the proportion of F1 seedlings with the respective genotypes if the mutant alleles promote no deleterious effects. A z-test for one proportion was performed to test if the observed homozygous mutant frequencies differ from the expected Mendelian value and the significance level was corrected for multiple comparisons using Bonferroni. Asterisks indicate significant differences from the expected value (25%). Primers used for genotyping are listed in Table S1.

| Genotype<br>(selfed) | Genotype of progeny |  |  | Number of<br>seeds (n) |
| --- | --- | --- | --- | --- |
|  | *cycb1;2<sup>+/+</sup>* | *cycb1;2<sup>+/-</sup>* | *cycb1;2<sup>-/-</sup>* |  |
| *cycb1;1<sup>-/-</sup> cycb1;2<sup>+/-</sup>* | 25.26% | 65.26% | 9.47%** | 95 |
| *gip1<sup>-/-</sup> cycb1;1<sup>-/-</sup>*<br>*cycb1;2<sup>+/-</sup>* | 31.25% | 64.58% | 4.17%**** | 96 |
| *gip2<sup>-/-</sup> cycb1;1<sup>-/-</sup>*<br>*cycb1;2<sup>+/-</sup>* | 34.55% | 65.44% | 0%**** | 191 |
| Expected Mendelian | 25% | 50% | 25% | - |

**Table 4.**
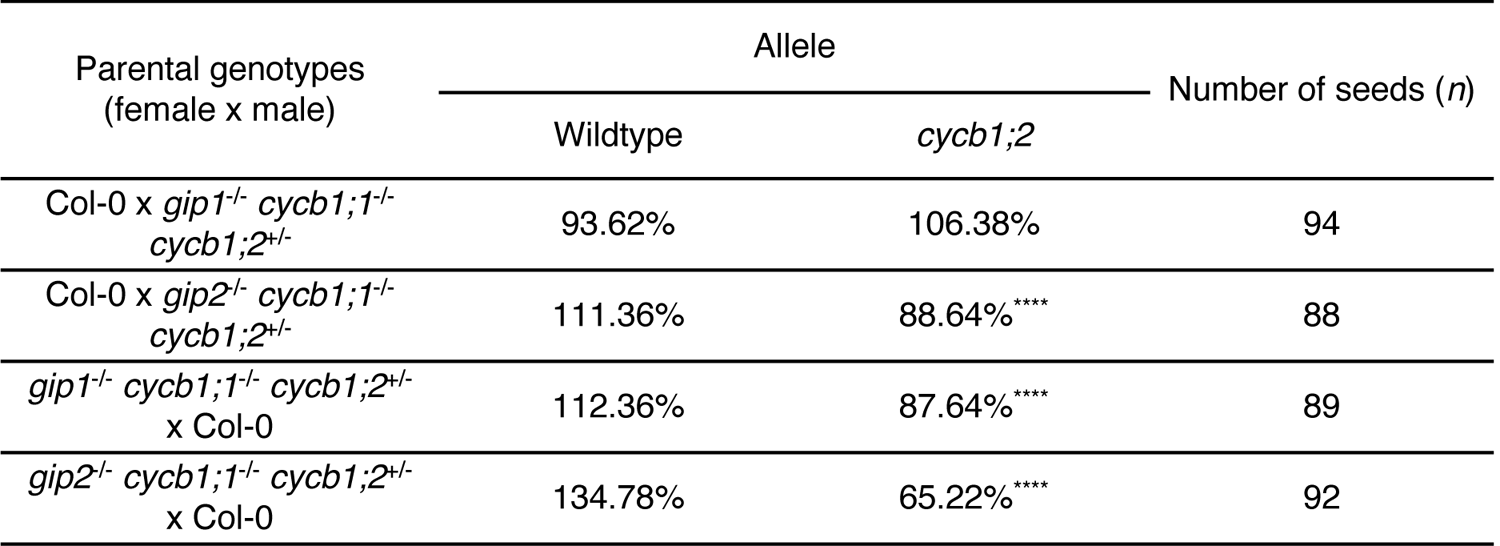
Transmission efficiency of the *cycb1;2* mutant allele in reciprocal crosses of ***gip1*^-/-^ *cycb1;1*^-/-^ *cycb1;2*^+/-^ and *gip2* ^-/-^ *cycb1;1*^-/-^ *cycb1;2*^+/-^ with wildtype.** A transmission efficiency of 100% indicates full transmission of the mutant allele, i.e. 50% of the genotyped F1 seedlings are heterozygous. A z-test for one proportion was performed to test if the observed transmission frequencies differ from expected values and the significance level was corrected for multiple comparisons using Bonferroni. Asterisks indicate significant differences from expected values. Primers used for genotyping are listed in Table S1.

Based on the results of our segregation analysis and reciprocal crosses (Tables 3 and 4), the assumption that GIP1 and GIP2 are completely interchangeable is challenged. It seems likely that GIP1 but not GIP2 is regulated by a CYCB1-dependent process. Thus, we generated a 2,849 bp genomic *GFP-GIP1* reporter in order to follow protein localization in the *cycb1;1 cycb1;2* mutant background (Fig 9). We reasoned that, if GIP1 is indeed modulated by CYCB1-CDK complexes, protein localization would be impaired in a *cycb1* mutant background.

**Figure 9.**
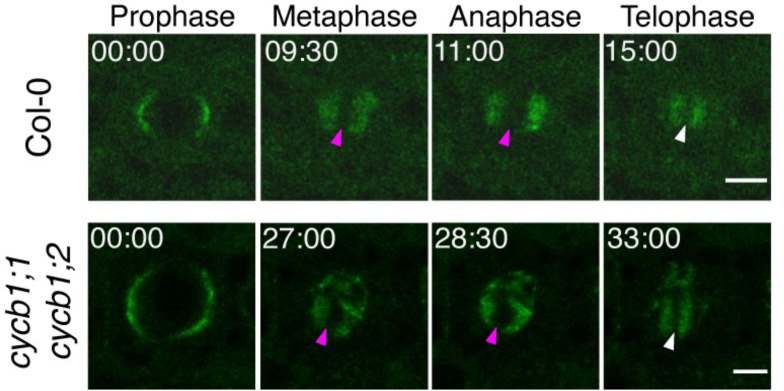
GIP1 mislocalizes in *cycb1;1 cycb1;2* mitosis. Time-lapse of confocal microscope pictures of root meristematic cells tagged with GFP-GIP1 in Col-0 (top panel) and *cycb1;1 cycb1;2* (bottom panel). GIP1 localizes at the nuclear polar caps, followed by co-localization with microtubules at the spindle and phragmoplast arrays. In *cycb1;1 cycb1;2* double mutants, GIP1 exhibited an abnormal localization, being found at the spindle (magenta arrowheads) and phragmoplast (white arrowheads) midzones, which are normally devoid of the protein. Scale bars: 5 μm.

GIP1 is a microtubule nucleation factor and mainly localizes to microtubule minus ends across mitosis. At prophase, it localizes at the nuclear surface. At metaphase and anaphase, it is directed to the two spindle poles, co-localizing with microtubule minus ends. At telophase, it localizes at the two opposing sides of the phragmoplast, directing microtubule nucleation towards the midzone. With some degree of variation between divisions, GFP-GIP1 localization differed greatly in *cycb1;1 cycb1;2* mutants in comparison to the wildtype. In some cases, GFP-GIP1 was found to remain in the spindle midzone (Fig 9; magenta arrowheads) during metaphase in abnormal mitotic divisions. The resulting phragmoplast, which is normally devoid of GIP1 in its midzone (Fig 9; white arrowheads), also contained remaining GIP1. The duration of these abnormal mitotic divisions in the double mutant was also around double the time of the wildtype divisions (Fig 9). Thus, we conclude that B1-type cyclins and in particular CYCB1;2 control microtubule organization through the regulation of GIP1 and likely several other substrates.

## Discussion

Angiosperms have undergone an extensive expansion of the cyclin family in comparison to yeast and mammals, containing for instance a total of 10 different cyclin families with more than 50 protein-encoding cyclin genes in Arabidopsis (Wang *et al*, 2004; Jia *et al*, 2014). For most of these genes, functional information is still lacking. Although genetic dissection of such an enlarged number of cyclin members may be more challenging and require the construction of multiple mutant combinations, it also provides an opportunity to study the function of these cyclins in compromised, yet viable mutant combinations of redundantly acting genes. In contrast, mutants for the CycB1 in mice, for example, are not viable and die *in utero* making its analysis, especially at the developmental level, challenging (Brandeis *et al*, 1998). Here, we have functionally dissected the group of B1-type cyclins and created various double and multiple mutant combinations. In particular the combination of *cycb1;1* and *cycb1;2* proved to be a valuable tool to study the function of this class of cyclins.

### CYCB1;2 is the central-most B1-type cyclin in *Arabidopsis*

When we analyzed the available information for the different *Arabidopsis* accessions using a public depository (TAIR), we found that very few SNPs are present in *CYCB1;2* and there was sequence information for the *CYCB1;2* coding region in the deposited genome data for all accessions. In contrast, *CYCB1;5* has accumulated different point mutations in the different *Arabidopsis* accessions, which supports our finding that *CYCB1;5* is likely a pseudogene. *CYCB1;1* and *CYCB1;3* displayed slightly more SNPs than *CYCB1;2*, which is in consistence with the overall normal growth and seed set of *cycb1;1 cycb1;3* mutants. In contrast, double mutants of either of them with *cycb1;2*, i.e., *cycb1;1 cycb1;2* and *cycb1;2 cycb1;3*, displayed strong mutant phenotypes. Thus, our work reveals a hierarchy of cyclin B1 function with CYCB1;2 being the most important mitotic regulator in this class, whose function is backed-up by CYCB1;1 and CYCB1;3 (Fig 10). CYCB1;4 comes at a third level that was found to act redundantly with CYCB1;1 and CYCB1;2 during female gametophyte development.

**Figure 10.**
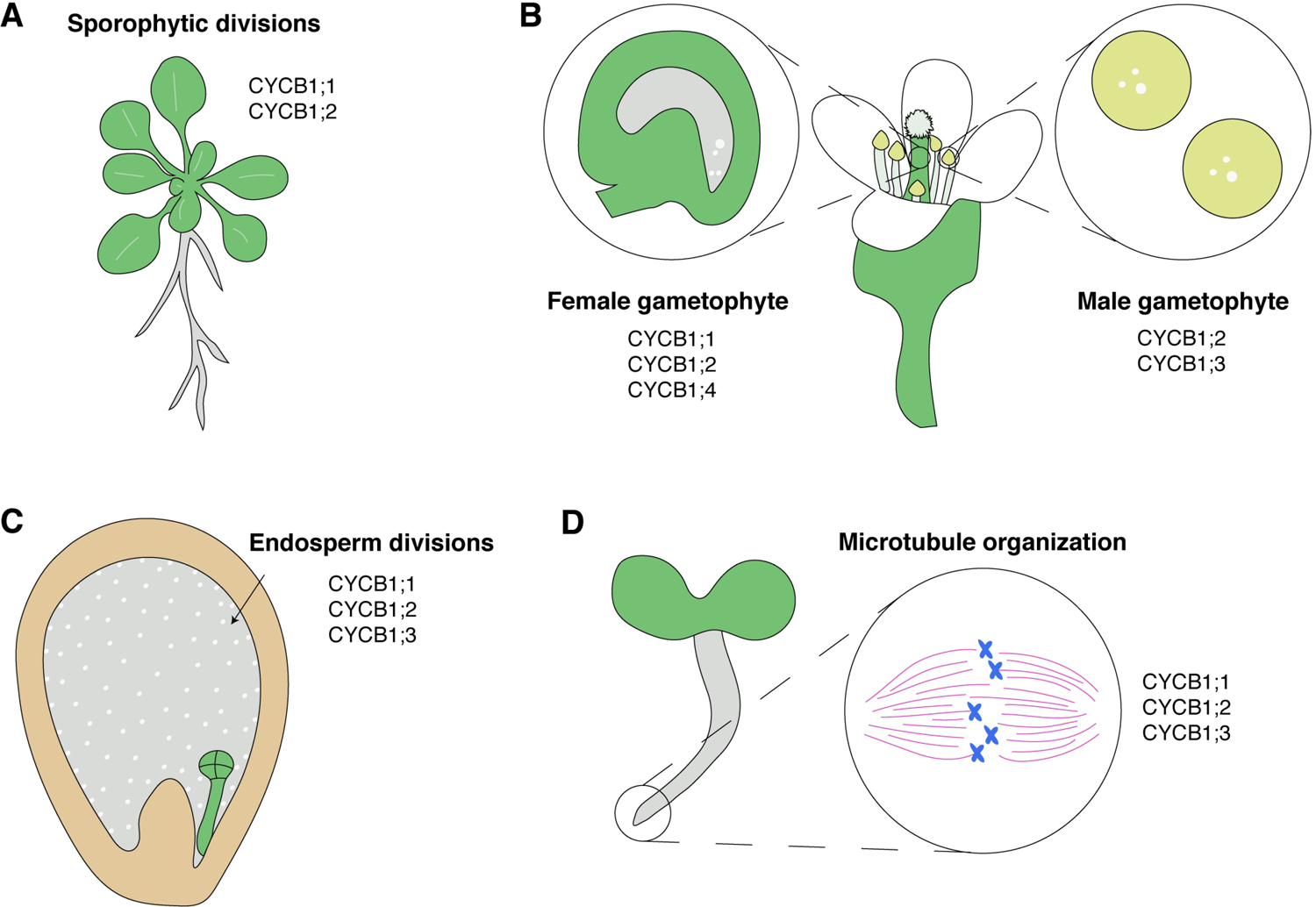
The differential role of the members of the CYCB1 group is developmental and tissue specific. A. CYCB1;1 and CYCB1;2 are the main B1-type cyclins involved in sporophytic divisions, as seen in the general dwarf phenotype of the *cycb1;1 cycb1;2*. B. CYCB1;1, CYCB1;2 and CYCB1;4 are mainly implicated in the female gametophyte development, as shown by the defects in embryo sacs of the triple mutants. CYCB1;2 and CYCB1;3 are primarily involved in the male gametophyte development, as seen by for instance the higher pollen death in mutant combinations of those two members. C. In the endosperm, the requirement of CYCB1;1, CYCB1;2 and CYCB1;3 for mitotic divisions became clear, as shown by the reduced proliferation of that tissue in double mutants of those cyclins. D. In mitotic divisions of the roots, microtubule arrays seem to be majorly regulated by CYCB1;1 and CYCB1;2, followed secondarily by CYCB1;3.

In parallel to a development-specific requirement, comes the role of B1-type cyclins in response to environmental condition. In particular, CYCB1;1 has been found to play a key role in homologous recombination repair during DNA damage and hence its upregulation can also indicate a cellular stress state (Weimer *et al*, 2016).

Thus, when describing cell proliferation in a developmental context and/or as response to environmental conditions, to mitotic gene to monitor needs to be carefully chosen, i.e. CYCB1;3 is not a good choice to describe female gametophyte development and CYCB1;1 has the ambiguity to monitor stress conditions. Due its central role, CYCB1;2 appears to be a good general choice, yet certain circumstances might make the analysis of other cyclin genes preferable.

### Endosperm – a demanding structure

During plant development, many different cell cycle programs are executed (Jakoby & Schnittger, 2004). One of the most particular proliferation modes are the free nuclear divisions during endosperm proliferation (Berger *et al*, 2006). Despite of its importance for seed growth and embryo nutrition, there is currently very little known about the cell cycle machinery that drives these free nuclear divisions. Laser-dissection microscopy-based transcriptional profiling of *Arabidopsis* endosperm revealed that B1-type cyclins are among the most prominently expressed cell-cycle regulators in this tissue (Day *et al*, 2008). Consistently, we found that nuclear divisions are reduced in *cycb1;2* single mutants and aberrant mitotic divisions appear in *cycb1;1 cycb1;2* and *cycb1;2 cycb1;3* double mutants. Correspondingly, seed development defects were reported as an effect of silencing cyclin B1 expression in rice (Guo *et al*, 2010).

Strikingly, the phenotypes of *cycb1;1 cycb1;2* and *cycb1;2 cycb1;3* double mutant endosperm closely resembles the defects seen in mutants for ENDOSPERM DEFECTIVE 1 (EDE1), a plant-specific microtubule binding protein (Pignocchi *et al*, 2009). EDE1 contains short CDK consensus phosphorylation sites (S/T-P) and so far has not been identified in CDK substrate searches in *Arabidopsis* (Pusch *et al*, 2012; Harashima *et al*, 2016). However, EDE1 could be phosphorylated by human Cdk complexes in *in vitro* kinase assays and it is known that short CDK consensus sites are sufficient to be phosphorylated by CDK/cyclin complexes (Pignocchi & Doonan, 2011; Ubersax *et al*, 2003). Interestingly, many cytoskeletal components are highly expressed in proliferating endosperm tissue and the free nuclear divisions might be very sensitive to alterations in cytoskeleton function, providing a possible reason why these divisions are apparently more sensitive to the loss of CYCB1 function (Day *et al*, 2009). Endosperm development in *Arabidopsis* might thus advance as a model system to study cell biological questions. However, endosperm is difficult to access, since it is buried in maternal structures, such as the seed coat and the silique. Therefore, morphological analyses always require mechanical preparation steps. In this light, the identification of homozygous *cycb1;1 cycb1;2* homozygous double mutants represents a unique tool to investigate the control of mitosis in roots or other much more easily accessible plant tissues than gametophytes.

### B1-type cyclins and the control of microtubule nucleation

Microtubules nucleate from ring-shaped complexes that contain γ-tubulin, and a family of related proteins called γ-tubulin complex proteins (GCPs). The composition of γ-tubulin ring complexes (γTURC) varies between organisms: budding yeast contains only the γ-tubulin small complex (γTUSC), with two molecules of γ-tubulin, and one each of GCP2 and GCP3 (Vinh *et al*, 2002). On the other hand, animal nucleating complexes are made of multiple copies of the γTUSC plus GCP4, GCP5 and GCP6, as well as other non-GCP constituents, such as GIP1/MOZART1, MOZART2 and NEDD1, which is a localization factor (Tovey & Conduit, 2018). In plants, γ-tubulin complexes contain all GCP subunits, the GIP1/MOZART1 protein, and a NEDD1 homolog (Lee & Liu, 2019).

The dynamic assembly and disassembly of the microtubule network generally runs in parallel with the cell cycle and, for example, even strong defects in microtubule arrays, e.g. lack of the PPB formation (Schaefer *et al*, 2017), do not block the cell cycle. However, rearrangements of the microtubule cytoskeleton in plant cells are obviously coupled with the cell cycle. Specific microtubule arrays accompany each stage of the cell cycle, either in interphase (the interphase cortical microtubule array), in pre-mitosis (the PPB), or in mitosis (spindle and phragmoplast). Moreover, several observations indicate a tight – at least temporal – coordination of both cycles. For instance, the PPB is formed in late G2-prophase in somatic tissues. Rapid PPB dismantling precisely coincides with nuclear envelope breakdown and entry into metaphase. Prospindle and spindle formation also take place at precise stages of the cell cycle. Likewise, the phragmoplast is precisely initiated at telophase from remnants of the spindle. However, very little is currently known about the molecular mechanisms of how this coordination is achieved. Interestingly, several cell cycle regulators including CDKA;1 have been identified at microtubule arrays such as the PPB, spindle and phragmoplast (Boruc *et al*, 2010). Our finding that GIP1/MOZART1 is phosphorylated by CDKB2-CYCB1 complexes offers a potential mechanism of how the cell cycle might orchestrate microtubule assembly. Interestingly, double PPBs and asymmetric PPBs, as we report here to be present in *cycb1;1 cycb1;2* mutants, have also been described in a *gip1 gip2* double knockdown mutant previously further strengthening that CYCB1 control the cytoskeleton via regulation of the γTURC complex (Janski *et al*, 2012).

Moreover, other factors of the γTURC have also been found to be phosphorylated in animals and yeast. For instance, all core units of the γTURC (γ-tubulin, GCP2-GCP6, GCP-WD and GCP8/MOZART2) but GIP1/MOZART1 have been found to be phosphorylated in mammals (Teixidó-Travesa *et al*, 2012). In particular, CDKs were shown to phosphorylate γTURC components including γ-tubulin and others in yeast and Nedd1 in humans (Zhang *et al*, 2009; Keck *et al*, 2011). However, the functional importance of these phosphorylation sites is not understood and an analysis of microtubule dynamics in animals is complicated due to the lethality of core cell cycle regulators such as Cdk1 or CycB1 (Santamaría *et al*, 2007; Brandeis *et al*, 1998).

In plants, all of the core γTURC components (GIP1, GCP2, GCP3, GCP4, GCP5a, GCP5b, NEDD1 and γ-tubulin 1 as well as γ-tubulin 2) but GIP2 have at least one CDK consensus phosphorylation site, and for NEDD1 and GCP4 as well as GCP5a a phosphorylated Ser/Thr in a consensus CDK site has been deposited in the PhosPhAt database (http://phosphat.uni-hohenheim.de). In addition, CYCB1;3 has been found to bind to GCP3 and γ-tubulin 1 (Van Leene *et al*, 2010). Thus, the regulation of the γTURC complex by CYCB1s likely goes even beyond the here reported phosphorylation of GIP1.

### The CYCB1 group has an evolutionarily conserved role in microtubule networks

Mammalian CycB1 is mainly cytoplasmic in interphase, rapidly accumulates in the nucleus at the end of prophase and associates with the mitotic apparatus in the course of mitosis, i.e., chromatin, microtubules, kinetochores and centrosomes (Hagting *et al*, 1999; Toyoshima *et al*, 1998; Yang *et al*, 1998; Bentley *et al*, 2007). Loss of the CycB1 function in mice results in very early embryo lethality (Brandeis *et al*, 1998). In contrast, mammalian cyclin B2 (CycB2) localizes mostly to the Golgi apparatus in both interphase and metaphase and CycB2 knock-out mice are viable (Brandeis *et al*, 1998; Jackman *et al*, 1995; Draviam *et al*, 2001). However, knocking down both CycB1 and CycB2 in HeLa cells showed a redundant function for both cyclins (Soni *et al*, 2008). Cyclin B3 is only poorly expressed in mitotic cells, but its mRNA is readily observed in both male and female meiosis (Nguyen *et al*, 2002; Lozano *et al*, 2002).

Interestingly, CycB1 in mammals localizes to the outer plate of the kinetochore at prometaphase and later on to the spindle poles following microtubule attachment to kinetochores (Chen *et al*, 2008; Bentley *et al*, 2007). Reduction of CycB1 by the use of RNA interference results in irregular attachment of kinetochores to microtubules, chromosome alignment defects and delays anaphase (Chen *et al*, 2008), which is reminiscent of the chromosome alignment and segregation problems in addition to the extended spindle stages we found in *cycb1;1 cycb1;2* mutants.

In contrast to many other eukaryotes, the setup of interphasic and mitotic microtubule networks in flowering plants is not driven by microtubule organizing centers containing centrioles/basal bodies. Instead, it has been proposed that mitotic microtubule networks nucleate from chromatin. Consistent with a role in microtubule nucleation, CYCB1;1 and CYCB1;2 have been found to be present mainly at chromatin during mitosis, while CYCB1;3 localized to both chromatin and cytoplasm and CYCB1;4 localized mainly in the cytoplasm as well as the region of the cytoplasm that co-localizes with the mitotic spindle (Bulankova *et al*, 2013). Thus, although the CYCB1 group in *Arabidopsis* appears from a general point of view to regulate the mitotic microtubule network similarly to CycB1 in mammals, the localization of B1-type cyclins is different in both species, indicating that the work of B-type cyclins in different species is differently distributed among its members.

Remarkably, and in contrast to CycB1 localization in mammals, CYCB1;1, CYCB1;2 and CYCB1;3 were not found at the mitotic spindle. We cannot rule out at the moment that B1-type cyclins do not have a function in further organizing the mitotic spindle. However, it seems likely that other, yet to be characterized subgroups of mitotic cyclins, in particular the B2- and B3 group, might play a key role here especially since a recent analysis of CYCB3;1 found that this cyclin is localized to the spindle, at least in meiosis (Sofroni *et al*, 2020). With this, it will be exciting to have eventually a complete view on B-type cyclin function in *Arabidopsis*.

## Materials and Methods

### Plant material and growth conditions

The accessions Columbia (Col-0) and Nossen (No-0) were used as wildtype controls. The single *cycb1;1*, *cycb1;2*, *cycb1*;3 and *cycb1;4* mutants were previously described and characterized (Weimer *et al*, 2016). The *cycb1;3* T-DNA insertion is in a No-0 background. The *gip1* (GABI_213D01) and *gip2* (FLAG_36406) mutants were also previously characterized (Janski *et al*, 2012). Genotyping primers are listed in Table S1.

*Arabidopsis thaliana* seeds were sown on half-strength (½) Murashige and Skoog (MS) medium (basal salt mixture, Duchefa Biochemie) containing 0.5% sucrose and 0.8% plant agar (Duchefa Biochemie) at pH 5.8. Seeds were either sterilized with chlorine gas or by liquid sterilization. For the liquid sterilization, a 2% bleach, 0.05% Triton X-100 solution was applied for 5 min, followed by three washing steps with sterile distilled water and the addition of 0.05% agarose. Stratification of the seeds on plates was performed at 4°C for 2 to 3 days in the dark. Plants were initially grown *in vitro* at 22°C in a 16-h light regime and subsequently transferred to soil with a 16-h light/21°C and 8-h/18°C dark regime with 60% humidity.

### Quantitative PCR

Total RNA was isolated from plant tissues using the RNeasy Plant Mini Kit (Qiagen). DNase (TaKaRa) treatment was performed to avoid DNA contamination and RNA concentration was measured using a Nanodrop ND-1000 instrument. 3.5 µg of total RNA was reverse transcribed using SuperScript® III reverse transcriptase kit (Invitrogen). An additional step of RNase H treatment at 37°C for 20 minutes was performed to eliminate remaining RNA. The cDNA was further purified and concentrated by using QIAquick PCR Purification Kit (Qiagen) and the concentration was determined by Nanodrop ND-1000 instrument. Finally, using cDNA as the template, qPCR was performed on a Light-cycler LC480 instrument (Roche) as per the manufacturer’s instructions.

### Plasmid construction and plant transformation

To analyze the expression of the *CYCB1* group in seeds, we used previously generated promoter reporter lines for *CYCB1;1* to *CYCB1;4* fused at the N-terminus to GFP (Weimer *et al*, 2016).

To generate a *PRO_GIP1_:GFP:GIP1* construct, a 2,849 bp genomic region including the native promoter and terminator was amplified by PCR and integrated into a *pENTR-D-TOPO* vector. A *SmaI* restriction site was introduced before the first *ATG* codon of the *GIP1* CDS. After linearization of the construct by restriction digest with *SmaI*, a ligation with *GFP* was performed, followed by LR reaction with the destination vector *pGWB501*. The constructs were transformed in *Arabidopsis thaliana* by floral dipping.

### Flow cytometry assay

Ten seven-day-old seedlings per genotype were chopped with a new razorblade in homogenization buffer (45 mM MgCl_2_, 20 mM MOPS, 30 mM sodium citrate, 0.1% Triton X-100, pH 7.0), followed by filtration through a 15-μm nylon mesh. After that, propidium iodide (Sigma) and RNase A (Sigma) were added to final concentrations of 50 μg/mL and 10 μg/mL respectively. Samples were left on ice for 5 min and then analyzed in a S3e Cell Sorter (Bio-Rad) with a laser excitation at 488 nm. The scatterplots were analyzed and processed using the FlowJo software.

### Endosperm nuclei proliferation analysis

Flower buds were initially emasculated before the visible maturation and release of pollen. Emasculated flowers were then hand-pollinated with pollen from the same genotype after 2 to 3 days. After 3 days, siliques were dissected and fixed in a solution of 4% glutaraldehyde in 12.5 mM cacodylate buffer, pH 6.8, followed by vacuum application for 20 min and storage at 4°C overnight. The following day, individual seeds were mounted on microscope slides containing a clearing 1:8:2 glycerol:chloral hydrate:water solution and stored at 4°C overnight. Imaging was performed with a Zeiss LSM 780 or 880 confocal microscope with excitation at 488 nm and detection between 498 and 586 nm and Z-stacks were analyzed using the Fiji software.

### Pollen staining

To identify single nuclei in mature pollen, pollen grains were released into a DAPI staining solution (2.5 μg/ml DAPI, 0.01% Tween, 5% dimethyl sulfoxide, 50 mM Na phosphate buffer, pH 7.2) and incubated at 4°C overnight. Pollen viability was analyzed by mounting pollen as previously described (Alexander, 1969). Imaging was performed with a Zeiss Axioimager.

### Embryo sac analysis

Mature ovules and developing seeds were prepared from siliques before and 3 days after fertilization, respectively, mounted on microscope slides on a clearing 1:8:2 glycerol:chloral hydrate:distilled water solution and kept at 4°C overnight before analysis as previously described (Nowack *et al*, 2006). Differential Interference Contrast (DIC) imaging was performed on a Zeiss Axioimager.

### Microtubule cytoskeleton dynamics in roots

Meristematic cell divisions in the root were observed in 5 to 7 day-old seedlings under a layer of ½ MS medium using a Leica TCS SP8 inverted confocal microscope.

### Immunostaining

Roots of 4-day-old Arabidopsis seedlings were fixed in 4% paraformaldehyde and 0.1% Triton X-100 in ½ MTSB buffer (25 mM PIPES, 2.5 mM MgSO_4_, 2.5 mM EGTA, pH 6.9) for 1 hour under vacuum, then rinsed in PBS 1X for 10 minutes. Samples were then permeabilized in ethanol for 10 minutes and rehydrated in PBS for 10 minutes. Cell walls were digested using the following buffer for one hour: 2 mM MES pH 5, 0.20% driselase and 0.15% macerozyme. Tissues were hybridized overnight at room temperature with the B-5-1-2 monoclonal anti-α-tubulin (Sigma) and the anti-KNOLLE antibody (kind gift of G. Jürgens, University of Tübingen, Germany; Lauber *et al*, 1997). The next day, tissues were washed for 15 minutes in PBS, 50 mM glycine, incubated with secondary antibodies (Alexa Fluor 555 goat anti-rabbit for KNOLLE antibody and Alexa Fluor 488 goat anti-mouse for the tubulin antibody) overnight and washed again in PBS, 50 mM glycine and DAPI 20 ng/ml. Tissues were mounted in VECTASHIELD and DAPI and viewed using an SP8 confocal laser microscope (Leica Microsystems). Samples were excited sequentially at 405 nm (DAPI), 488 nm (@TUB/Alexa Fluor 488), and 561 nm (@KNOLLE/Alexa Fluor 555), with an emission band of 420-450 nm (DAPI), 495-545 nm (Alexa Fluor 488), and 560-610 nm (Alexa Fluor 555) using a PMT for DAPI imaging, and hybrid detectors for MT and KNOLLE imaging. All stacks have been imaged using the same zoom (x 1.60) with a pixel size xyz of 200 nm x 200 nm x 500 nm.

A blind counting was set up to count mitotic MT arrays seen in ten roots of each genotype. The “Cell counter” plugin was used to count the occurrence of MT arrays within each root stack (https://imagej.nih.gov/ij/plugins/cell-counter.html)

### Protein expression and purification and *in vitro* kinase assays

Purified GIP1 was kindly donated by Nicolas Baumberger (IBMP, Strasbourg). *GIP1* CDS was cloned into the *pGEX-2TK* vector (GE Healthcare; courtesy of Etienne Herzog) and transformed into the BL21(DE3) *E. coli* strain. An overnight culture cultivated at 37°C was used to inoculate an expression culture at an OD_600_ of 0.1. The expression culture was grown at 37°C and 250 rpm until it reached an OD_600_ of 0.6. Afterwards, 0.5mM IPTG was added and the growth continued at 37°C for 6 hours. Cells were collected by centrifugation at 5000 x*g* for 15 min and the pellet resuspended in 50 mM Tris pH 8, 300 mM NaCl, 5% glycerol, 5 mM EDTA, 0.1% Tween 20. Cells were lyzed by sonication and the lysate clarified by centrifugation at 10,000 x*g* for 20 min at 4°C. GST-GIP was purified by passage onto a glutathione-sepharose GSTrap HP 1ml column (GE Healthcare) with 50 mM Tris pH 7.5, 10 mM MgCl2, 100 mM NaCl as equilibration/washing buffer and 50 mM Tris pH 7.5, 10 mM MgCl2, 100 mM NaCl plus 10 mM reduced glutathione as elution buffer. Elution fractions were analyzed on polyacrylamide gel and concentrated by ultrafiltration before being frozen and stored at −80°C. Histone H1^0^ was purchased from NEB. *In vitro* kinase assays were performed as described previously (Harashima & Schnittger, 2012).

### Oryzalin root growth assays

Plants were sown on ½ MS medium containing either 0.05% DMSO as a control or oryzalin. 100 mM oryzalin stock solutions were prepared in DMSO and stored at −20°C and further diluted to a final concentration of 150 nM or 200 nM for the root assays. Root growth was recorded daily up until 5 days after germination, when plates were scanned, and root length was subsequently measured using the Fiji software. Three biological replicates with at least 10 plants per genotype were performed. The mean root length of each individual experiment was determined and again averaged.

## Statistical analysis

The employed statistical tests are indicated in the figure legends. Statistical tests were performed using the GraphPad Prism 9 software and the XLSTAT plugin for Microsoft Excel. The distribution of the measured values was tested beforehand, e.g. by the Anderson-Darling test. If the distribution was significantly different from a normal distribution, a non-parametric test was employed. Significance levels: *P* ≥ 0.05 (ns), *P* < 0.05 (*), *P* < 0.01 (**), *P* < 0.001 (***), *P* < 0.0001 (****). In the case of the Chi-squared test followed by the Marascuilo procedure, significant pairwise comparisons are indicated by letters.

## Acknowledgements

This work was supported by a grant from the Ohsumi Frontier Science Foundation to S.K., by the Ministry of Education, Youth, and Sports of the Czech Republic, European Regional Development Fund-Project REMAP (grant CZ.02.1.01/0.0/0.0/15_003/0000479) to K.R., and by the Human Frontier Science Program to D.B. and A.S.

## Author contributions

MM, XZ, MP, FR, HH, SK, MH and AS designed the research; MM, XZ, MP, KB, FR, MK, HH, SK, and PB performed the experiments; MM, XZ, and MP performed the statistical analysis; MM, XZ, MP, KB, FR, MK, HH, SK, PB, MH, KR, DB, and AS analyzed and discussed the data; DB, KR and AS provided material and reagents, MM and AS wrote the article; MM, XZ, MP, FR, MK, HH, SK, PB, MH, KR, DB, AS revised and approved the article.

## Expanded View Figure legends

Figure EV1. *CYCB1;5* is a pseudogene. A-C. The predicted gene structure of *CYCB1;5*, including the observed cDNAs with exon skipping (A), intron retention and premature polyadenylation sites (B), and alternative splicing of a large intron (C).

Figure EV2. *CYCB1;1*, *CYCB1;2* and *CYCB1;3* but not *CYCB1;4* are expressed during seed development. A-C. Confocal microscope pictures of seeds expressing either *proCYCB1;1:GFP*, *proCYCB1;2:GFP* or *proCYCB1;3:GFP* in Col-0 (A), *cycb1;1 cycb1;2* (B) or *cycb1;2 cycb1;3* (C). Scale bars: 30 μm.

Figure EV3. Oryzalin root growth assays across time in *cycb1* single mutants. A-C. Quantification of root length in a control condition (A), 150 nM oryzalin (B) and 200 nM oryzalin (C). Graphs show mean ± SD of three biological replicates with at least 10 plants per genotype per replicate. Asterisks indicate a significant difference in root length in a two-way ANOVA followed by Tukey’s multiple comparisons test.

Figure EV4. Oryzalin root growth assays across time in *cycb1* double mutants. A-C. Quantification of root length in a control condition (A), 150 nM oryzalin (B) and 200 nM oryzalin (C). Graphs show mean ± SD of three biological replicates with at least 10 plants per genotype per replicate. Asterisks indicate a significant difference in root length in a two-way ANOVA followed by Tukey’s multiple comparisons test.

Figure EV5. **Ploidy analysis of young seedlings of the single and double mutants.** Flow cytometrical quantification of the different nuclear ploidies, as indicated by propidium iodide (PI) intensity.

## Data availability

This study includes no data deposited in external repositories.

## References

1. Alexander MP (1969) Differential staining of aborted and nonaborted pollen. Biotech Histochem 44: 117–122

2. Bentley AM, Normand G, Hoyt J & King RW (2007) Distinct Sequence Elements of Cyclin B1 Promote Localization to Chromatin, Centrosomes, and Kinetochores during Mitosis. Mol Biol Cell 18: 4847–4858

3. Berger F, Grini PE & Schnittger A (2006) Endosperm: an integrator of seed growth and development. Curr Opin Plant Biol 9: 664–670

4. Blethrow JD, Glavy JS, Morgan DO & Shokat KM (2008) Covalent capture of kinase-specific phosphopeptides reveals Cdk1-cyclin B substrates. Proc Natl Acad Sci U S A 105: 1442–1447

5. Boisnard-Lorig C, Colon-Carmona A, Bauch M, Hodge S, Doerner P, Bancharel E, Dumas C, Haseloff J & Berger F (2001) Dynamic analyses of the expression of the histone::YFP fusion protein in Arabidopsis show that syncytial endosperm is divided in mitotic domains. Plant Cell 13: 495–509

6. Boruc J, Mylle E, Duda M, de Clercq R, Rombauts S, Geelen D, Hilson P, Inzé D, van Damme D & Russinova E (2010) Systematic localization of the Arabidopsis core cell cycle proteins reveals novel cell division complexes. Plant Physiol 152: 553–565

7. Boudolf V, Lammens T, Boruc J, van Leene J, van den Daele H, Maes S, van Isterdael G, Russinova E, Kondorosi E, Witters E, et al (2009) CDKB1;1 forms a functional complex with CYCA2;3 to suppress endocycle onset. Plant Physiol 150: 1482–1493

8. Brandeis M, Rosewell I, Carrington M, Crompton T, Jacobs MA, Kirk J, Gannon J & Hunt T (1998) Cyclin B2-null mice develop normally and are fertile whereas cyclin B1-null mice die in utero. Proc Natl Acad Sci U S A 95: 4344–4349

9. Bulankova P, Akimcheva S, Fellner N & Riha K (2013) Identification of Arabidopsis Meiotic Cyclins Reveals Functional Diversification among Plant Cyclin Genes. PLoS Genet 9

10. Chen Q, Zhang X, Jiang Q, Clarke PR & Zhang C (2008) Cyclin B1 is localized to unattached kinetochores and contributes to efficient microtubule attachment and proper chromosome alignment during mitosis. Cell Res 18: 268–280

11. Cools T, Iantcheva A, Weimer AK, Boens S, Takahashi N, Maes S, van den Daele H, van Isterdael G, Schnittger A & de Veylder L (2011) The Arabidopsis thaliana checkpoint kinase WEE1 protects against premature vascular differentiation during replication stress. Plant Cell 23: 1435–1448

12. Day RC, Herridge RP, Ambrose BA & Macknight RC (2008) Transcriptome analysis of proliferating Arabidopsis endosperm reveals biological implications for the control of syncytial division, cytokinin signaling, and gene expression regulation. Plant Physiol 148: 1964–1984

13. Day RC, Müller S & Macknight RC (2009) Identification of cytoskeleton-associated genes expressed during Arabidopsis syncytial endosperm development. Plant Signal Behav 4: 883–886

14. Dewitte W, Scofield S, Alcasabas AA, Maughan SC, Menges M, Braun N, Collins C, Nieuwland J, Prinsen E, Sundaresan V, et al (2007) Arabidopsis CYCD3 D-type cyclins link cell proliferation and endocycles and are rate-limiting for cytokinin responses. Proc Natl Acad Sci U S A 104: 14537–14542

15. Dissmeyer N, Weimer AK, Pusch S, de Schutter K, Kamei CLA, Nowack MK, Novak B, Duan GL, Zhu YG, de Veylder L, et al (2009) Control of cell proliferation, organ growth, and DNA damage response operate independently of dephosphorylation of the arabidopsis Cdk1 Homolog CDKA;1. Plant Cell 21: 3641–3654

16. Dissmeyer N, Weimer AK, de Veylder L, Novak B & Schnittger A (2010) The regulatory network of cell cycle progression is fundamentally different in plants versus yeast or metazoans. Plant Signal Behav 5

17. Doerner P, Jørgenser JE, You R, Steppuhn J & Lamb C (1996) Control of root growth and development by cyclin expression. Nature 380: 520–523

18. Draviam VM, Orrechia S, Lowe M, Pardi R & Pines J (2001) The localization of human cyclins B1 and B2 determines CDK1 substrate specificity and neither enzyme requires MEK to disassemble the Golgi apparatus. J Cell Biol 152: 945–958

19. Drews GN & Yadegari R (2002) Development and function of the angiosperm female gametophyte. Annu Rev Genet 36: 99–124

20. Furuno N, Elzen N Den & Pines J (1999) Human cyclin A is required for mitosis until mid prophase. J Cell Biol 147: 295–306

21. Guo J, Wang F, Song J, Sun W & Zhang XS (2010) The expression of Orysa;CycB1;1 is essential for endosperm formation and causes embryo enlargement in rice. Planta 231: 293–303

22. Haase SB, Winey M & Reed SI (2001) Multi-step control of spindle pole body duplication by cyclin-dependent kinase. Nat Cell Biol 3: 38–42

23. Hagting A, Jackman M, Simpson K & Pines J (1999) Translocation of cyclin B1 to the nucleus at prophase requires a phosphorylation-dependent nuclear import signal. Curr Biol 9: 680–689

24. Hagting A, Karlsson C, Clute P, Jackman M & Pines J (1998) MPF localization is controlled by nuclear export. EMBO J 17: 4127–4138

25. Harashima H, Dissmeyer N, Hammann P, Nomura Y, Kramer K, Nakagami H & Schnittger A (2016) Modulation of plant growth in vivo and identification of kinase substrates using an analog-sensitive variant of CYCLIN-DEPENDENT KINASE A;1. BMC Plant Biol 16: 1–19

26. Harashima H & Schnittger A (2012) Robust reconstitution of active cell-cycle control complexes from co-expressed proteins in bacteria. Plant Methods 8: 1–9

27. Jackman M, Firth M & Pines J (1995) Human cyclins B1 and B2 are localized to strikingly different structures: B1 to microtubules, B2 primarily to the Golgi apparatus. EMBO J 14: 1646–1654

28. Jakoby M & Schnittger A (2004) Cell cycle and differentiation. Curr Opin Plant Biol 7: 661– 669

29. Janski N, Masoud K, Batzenschlager M, Herzog E, Evrard JL, Houlné G, Bourge M, Chabouté ME & Schmit AC (2012) The GCP3-interacting proteins GIP1 and GIP2 are required for γ-tubulin complex protein localization, spindle integrity, and chromosomal stability. Plant Cell 24: 1171–1187

30. Jia RD, Guo CC, Xu GX, Shan HY & Kong HZ (2014) Evolution of the cyclin gene family in plants. J Syst Evol 52: 651–659

31. Keck JM, Jones MH, Wong CCL, Binkley J, Chen D, Jaspersen SL, Holinger EP, Xu T, Niepel M, Rout MP, et al (2011) A Cell Cycle Phosphoproteome of the Yeast Centrosome. 332: 1557–1562

32. Lacey KR, Jackson PK & Stearns T (1999) Cyclin-dependent kinase control of centrosome duplication. Proc Natl Acad Sci U S A 96: 2817–2822

33. Lauber MH, Waizenegger I, Steinmann T, Schwarz H, Mayer U, Hwang I, Lukowitz W & Jürgens G (1997) The Arabidopsis KNOLLE protein is a cytokinesis-specific syntaxin. J Cell Biol 139: 1485–1493

34. Lee YRJ & Liu B (2019) Microtubule nucleation for the assembly of acentrosomal microtubule arrays in plant cells. New Phytol 222: 1705–1718

35. Van Leene J, Hollunder J, Eeckhout D, Persiau G, Van De Slijke E, Stals H, Van Isterdael G, Verkest A, Neirynck S, Buffel Y, et al (2010) Targeted interactomics reveals a complex core cell cycle machinery in Arabidopsis thaliana. Mol Syst Biol 6

36. Lindqvist A, Rodríguez-Bravo V & Medema RH (2009) The decision to enter mitosis: feedback and redundancy in the mitotic entry network. J Cell Biol 185: 193–202

37. Lozano JC, Schatt P, Peaucellier G, Picard A, Perret E & Arnould C (2002) Molecular cloning, gene localization, and structure of human cyclin B3. Biochem Biophys Res Commun 291: 406–413

38. McCormick S (2004) Control of male gametophyte development. Plant Cell 16: 142–154

39. Menges M, De Jager SM, Gruissem W & Murray JAH (2005) Global analysis of the core cell cycle regulators of Arabidopsis identifies novel genes, reveals multiple and highly specific profiles of expression and provides a coherent model for plant cell cycle control. Plant J 41: 546–566

40. Morgan DO (1997) CYCLIN-DEPENDENT KINASES : Engines, Clocks, and Microprocessors. Annu Rev Cell Dev Biol 13: 261–291

41. Nakamura M, Yagi N, Kato T, Fujita S, Kawashima N, Ehrhardt DW & Hashimoto T (2012) Arabidopsis GCP3-interacting protein 1/MOZART 1 is an integral component of the γ-tubulin-containing microtubule nucleating complex. Plant J 71: 216–225

42. Nakayama KI & Nakayama K (2005) Regulation of the cell cycle by SCF-type ubiquitin ligases. Semin Cell Dev Biol 16: 323–333

43. Nguyen TB, Manova K, Capodieci P, Lindon C, Bottega S, Wang XY, Refik-Rogers J, Pines J, Wolgemuth DJ & Koff A (2002) Characterization and expression of mammalian cyclin B3, a prepachytene meiotic cyclin. J Biol Chem 277: 41960–41969

44. Nowack MK, Grini PE, Jakoby MJ, Lafos M, Koncz C & Schnittger A (2006) A positive signal from the fertilization of the egg cell sets off endosperm proliferation in angiosperm embryogenesis. Nat Genet 38: 63–67

45. O’Farrell PH (2001) Triggering the all-or-nothing switch into mitosis. Trends Cell Biol 11: 512–519

46. Pastuglia M, Azimzadeh J, Goussot M, Camilleri C, Belcram K, Evrard JL, Schmit AC, Guerche P & Bouchez D (2006) γ-tubulin is essential for microtubule organization and development of Arabidopsis. Plant Cell 18: 1412–1425

47. Pignocchi C & Doonan JH (2011) Interaction of a 14-3-3 protein with the plant microtubule-associated protein EDE1. Ann Bot 107: 1103–1109

48. Pignocchi C, Minns GE, Nesi N, Koumproglou R, Kitsios G, Benning C, Lloyd CW, Doonan JH & Hills MJ (2009) Endosperm Defective1 is a novel microtubule-associated protein essential for seed development in Arabidopsis. Plant Cell 21: 90–105

49. Pusch S, Harashima H & Schnittger A (2012) Identification of kinase substrates by bimolecular complementation assays. Plant J 70: 348–356

50. Riabowol K, Draetta G, Brizuela L, Vandre D & Beach D (1989) The cdc2 kinase is a nuclear protein that is essential for mitosis in mammalian cells. Cell 57: 393–401

51. Santamaría D, Barrière C, Cerqueira A, Hunt S, Tardy C, Newton K, Cáceres JF, Dubus P, Malumbres M & Barbacid M (2007) Cdk1 is sufficient to drive the mammalian cell cycle. Nature 448: 811–815

52. Schaefer E, Belcram K, Uyttewaal M, Duroc Y, Goussot M, Legland D, Laruelle E, De Tauzia-Moreau ML, Pastuglia M & Bouchez D (2017) The preprophase band of microtubules controls the robustness of division orientation in plants. Science (80-) 356: 186–189

53. Schnittger A, Schöbinger U, Bouyer D, Weinl C, Stierhof YD & Hülskamp M (2002) Ectopic D-type cyclin expression induces not only DNA replication but also cell division in Arabidopsis trichomes. Proc Natl Acad Sci U S A 99: 6410–6415

54. De Schutter K, Joubès J, Cools T, Verkest A, Corellou F, Babiychuk E, Van Der Schueren E, Beeckman T, Kushnir ST, Inzé D, et al (2007) Arabidopsis WEE1 kinase controls cell cycle arrest in response to activation of the DNA integrity checkpoint. Plant Cell 19: 211–225

55. Sofroni K, Takatsuka H, Yang C, Dissmeyer N, Komaki S, Hamamura Y, Böttger L, Umeda M & Schnittger A (2020) CDKD-dependent activation of CDKA;1 controls microtubule dynamics and cytokinesis during meiosis. J Cell Biol 219: 1–21

56. Soni D V., Sramkoski RM, Lam M, Stefan T & Jacobberger JW (2008) Cyclin B1 is rate limiting but not essential for mitotic entry and progression in mammalian somatic cells. Cell Cycle 7: 1285–1300

57. Teixidó-Travesa N, Roig J & Lüders J (2012) The where, when and how of microtubule nucleation - one ring to rule them all. J Cell Sci 125: 4445–4456

58. Tovey CA & Conduit PT (2018) Microtubule nucleation by γ-tubulin complexes and beyond. Essays Biochem 62: 765–780

59. Toyoshima F, Moriguchi T, Wada A, Fukuda M & Nishida E (1998) Nuclear export of cyclin B1 and its possible role in the DNA damage-induced G2 checkpoint. EMBO J 17: 2728–2735

60. Tyson JJ & Novak B (2001) Regulation of the eukaryotic cell cycle: Molecular antagonism, hysteresis, and irreversible transitions. J Theor Biol 210: 249–263

61. Ubersax JA, Woodbury EL, Quang PN, Paraz M, Blethrow JD, Shah K, Shokat KM & Morgan DO (2003) Targets of the cyclin-dependent kinase Cdk1. Nature 425: 859–864

62. Vandepoele K, Raes J, De Veylder L, Rouzé P, Rombauts S & Inzé D (2002) Genome-wide analysis of core cell cycle genes in Arabidopsis. Plant Cell 14: 903–916

63. Vanneste S, Coppens F, Lee E, Donner TJ, Xie Z, Van Isterdael G, Dhondt S, De Winter F, De Rybel B, Vuylsteke M, et al (2011) Developmental regulation of CYCA2s contributes to tissue-specific proliferation in Arabidopsis. EMBO J 30: 3430–3441

64. Vinh DBN, Kern JW, Hancock WO, Howard J & Davis TN (2002) Reconstitution and Characterization of Budding Yeast γ-Tubulin Complex. Mol Biol Cell 13: 1144–1157

65. Wang G, Kong H, Sun Y, Zhang X, Zhang W, Altman N, DePamphilis CW & Ma H (2004) Genome-wide analysis of the cyclin family in arabidopsis and comparative phylogenetic analysis of plant cyclin-like proteins. Plant Physiol 135: 1084–1099

66. Weimer AK, Biedermann S, Harashima H, Roodbarkelari F, Takahashi N, Foreman J, Guan Y, Pochon G, Heese M, Van Damme D, et al (2016) The plant-specific CDKB 1-CYCB 1 complex mediates homologous recombination repair in Arabidopsis. EMBO J 35: 2068–2086

67. Yamada M & Goshima G (2017) Mitotic spindle assembly in land plants: Molecules and mechanisms. Biology (Basel) 6

68. Yang J, Bardes ESG, Moore JD, Brennan J, Powers MA & Kornbluth S (1998) Control of Cyclin B1 localization through regulated binding of the nuclear export factor CRM1. Genes Dev 12: 2131–2143

69. Zhang X, Chen Q, Feng J, Hou J, Yang F, Liu J, Jiang Q & Zhang C (2009) Sequential phosphorylation of Nedd1 by Cdk1 and Plk1 is required for targeting of the γTuRC to the centrosome. J Cell Sci 122: 2240–2251

